# Information theory for data-driven model reduction in physics and biology

**DOI:** 10.1101/2024.04.19.590281

**Authors:** Matthew S. Schmitt, Maciej Koch-Janusz, Michel Fruchart, Daniel S. Seara, Michael Rust, Vincenzo Vitelli

## Abstract

Model reduction is the construction of simple yet predictive descriptions of the dynamics of many-body systems in terms of a few relevant variables. A prerequisite to model reduction is the identification of these relevant variables, a task for which no general method exists. Here, we develop a systematic approach based on the information bottleneck to identify the relevant variables, defined as those most predictive of the future. We elucidate analytically the relation between these relevant variables and the eigenfunctions of the transfer operator describing the dynamics. Further, we show that in the limit of high compression, the relevant variables are directly determined by the slowest-decaying eigenfunctions. Our information-based approach indicates when to optimally stop increasing the complexity of the reduced model. Furthermore, it provides a firm foundation to construct interpretable deep learning tools that perform model reduction. We illustrate how these tools work in practice by considering uncurated videos of atmospheric flows from which our algorithms automatically extract the dominant slow collective variables, as well as experimental videos of cyanobacteria colonies in which we discover an emergent synchronization order parameter.

**Significance Statement:** The first step to understand natural phenomena is to intuit which variables best describe them. An ambitious goal of artificial intelligence is to automate this process. Here, we develop a framework to identify these relevant variables directly from complex datasets. Very much like MP3 compression is about retaining information that matters most to the human ear, our approach is about keeping information that matters most to predict the future. We formalize this insight mathematically and systematically answer the question of when to stop increasing the complexity of minimal models. We illustrate how interpretable deep learning tools built on these ideas reveal emergent collective variables in settings ranging from satellite recordings of atmospheric fluid flows to experimental videos of cyanobacteria colonies.

---

The exhaustive description of a biological or physical system is usually impractical due to the sheer volume of information involved. As an example, the air in your office may be described by a 10^27^–dimensional state vector containing the positions and momenta of every particle in the room. Yet, for most practical purposes, it can be effectively described using only a small number of quantities such as pressure and temperature. Similar reductions can be achieved for systems ranging from diffusing particles to biochemical molecules and complex networks. In all cases, certain *relevant variables* can be predicted far into the future even though individual degrees of freedom in the system are effectively unpredictable.

The process by which one goes from the complete description of a system to a simpler one is known as model reduction. Diverse procedures for model reduction exist across the natural sciences. They range from analytical methods, such as adiabatic elimination and multiple-scale analysis (1–9), to data-driven methods such as independent component analysis (10), dynamic mode decomposition (11–13), diffusion maps (14), spectral submanifolds (15, 16), and deep encoder-decoder neural networks (17–24).

The success of these approaches is limited by a fundamental difficulty: before performing any reduction, one has to identify a decomposition of the full system into relevant and irrelevant variables. In the absence of prior knowledge and intuition (e.g. a clear separation of scales), identifying such a decomposition is an open problem (4). It may not even be clear *a priori* when to stop increasing the complexity of a simplified model or, conversely, when to stop reducing the amount of information needed to represent the dynamical state of a complex system. In both cases one must first determine the minimal number of relevant variables that are needed. The answer to this question depends in fact on how precisely and how far in the future you wish to forecast. Nonetheless, this answer should be compatible with fundamental constraints on forecasting set by external perturbations and finite measurement accuracy (25, 26).

In order to address the difficulty identified in the previous paragraph, we develop an information-theoretic framework for model reduction. Very much like MP3 compression is about retaining information that matters most to the human ear (27), model reduction is about keeping information that matters most to predict the future (28, 29). Inspired by this simple insight, we formalize model reduction as a lossy compression problem known as the information bottleneck (IB) **?**(30, 31). This formal step allows us to give a precise answer to the question of how to identify relevant and irrelevant variables. We show how and under what conditions the standard operator-theoretic formalism of dynamical systems (19, 32), which underlies most methods of model reduction, naturally emerges from optimal compression. Crucially, our framework systematically answers the question of when to stop increasing the complexity of a minimal model. Further, it provides a firm foundation to address a practical problem: the construction of deep learning tools to perform model reduction that are guaranteed to be interpretable. We illustrate our approach on benchmark dynamical systems and demonstrate that it works even on uncurated datasets, such as satellite movies of atmospheric flows downloaded directly from YouTube and biological datasets composed of microscopy videos of cyanobacteria colonies in which we discover an emergent synchronization order parameter.

## 1. Model reduction as a compression problem

We present here a method to extract collective variables most predictive of the system’s future evolution directly from data. This data is composed of a time sequence of measured states *x*_1_, *x*_2_, …, *x*_*T*_. The system state *x*_*t*_ could correspond to anything from the position of a single particle to an image of a fluid flow or the fluorescent molecules in a living system (Fig. 1a). The full state can be very high dimensional, with a number of dimensions equal to the number of observed pixels in the case of imaging data. However, the variation of any individual pixel is often of limited interest to us, as noise (either inherent or due to measurement) induces uncertainty about its true value. Individual pixels are, due to this uncertainty, poor predictors of the future state of the system. We can say that they are *irrelevant* for predicting the future. On the other hand, certain spatially-averaged collective variables may evolve slowly in time, and the future state of the system may be reliably estimated from them.

**Fig. 1.**
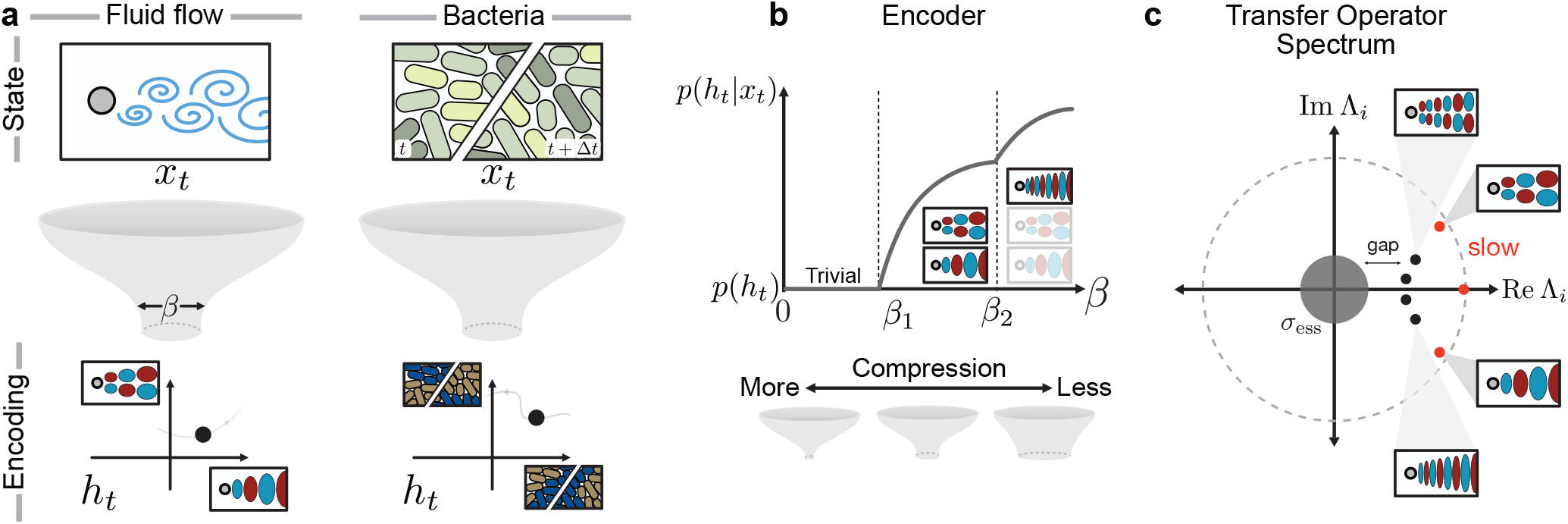
Interpretable dynamical variables in reduced models via the information bottleneck. (a) The information bottleneck compresses high-dimensional state variables *x*_*t*_, into simpler encoding variables *h*_*t*_ providing a controllable trade-off between the degree of compression and the predictive power about the system’s future. With deep neural networks, the encoding can be computed directly from data of observed fluid flows (left) or biological datasets, such as fluorescently labeled bacteria colonies (right). In general, the state of the variable *x*_*t*_ may comprise time-lagged variables of the intensity field, *x*_*t*_ = {*I*_*t*_, *I*_*t*+Δ*t*_} (right). The amount of compression is determined by the “width” of the bottleneck *β* [see Eq. (1))]. The resulting compressed, or encoded, variables *h*_*t*_ represent collective variables most predictive of the system’s future. (b) Schematic evolution of the encoder *p*(*h*_*t*_|*x*_*t*_) for varying compression strength *β*. For low *β* (high compression), the encoder is trivial and forgets everything about the input *x*_*t*_. After the first IB transition at *β*_1_, the encoder becomes non-trivial by gaining some dependence on *x*_*t*_; some features of the input are able to pass through the bottleneck. At subsequent IB transitions, additional features are learned. (c) The point spectrum of the transfer operator contains several slowly decaying modes (red). We show that the most predictive variables that IB systematically extracts correspond to the slowest eigenfunctions of the transfer operator, associated to eigenvalues Λ_*i*_ with |Λ_*i*_| ≈ 1. In fluid flows, the slowest-decaying eigenfunctions typically represent large-scale coherent patterns of the flow field, while faster-decaying eigenfunctions correspond to variations over shorter length scales.

We seek a way to “encode” each state *x*_*t*_ into a simple, lower-dimensional representation *h*_*t*_ in a way that isolates these relevant features of the input *x*_*t*_. For instance, in Fig. 1a, both the velocity field of a fluid flow and the images of a dynamic cyanobacteria colony (upper row) may be encoded as a point in a 2D space (lower row). The encoding is given by a probabilistic mapping *p*(*h*_*t*_|*x*_*t*_) which can be thought of as a machine which takes a state *x*_*t*_ and assigns it to a value *h*_*t*_. The fact that this mapping is probabilistic simply means we may have some uncertainty about the true value of *h*_*t*_ even given a measurement of the state. Whether or not some encoding is extracting “relevant” features of the state *x*_*t*_ is determined by the extent to which we can use it to predict the future. This predictive power can be quantified by the *mutual information* between the encoding and the future state, *I*(*H*_*t*_, *X*_*t*+Δ*t*_), which tells us how much the knowledge of *H*_*t*_ reduces our uncertainty about the future *X*_*t*+Δ*t*_ (see Methods). (In our notation, upper-case *X*_*t*_ refers to the random variable, while *x*_*t*_ refers to a particular value taken by the random variable.) To find a good encoder that extracts relevant features, we might try to find an encoding which maximizes this information. However, an encoder obtained in this way would simply copy the original state, *h*_*t*_ = *x*_*t*_, since *x*_*t*_ represents all the information we have about the system. In order to encourage the encoder to discard irrelevant features, we simultaneously seek an encoder which maximizes compression by minimizing the information about the original state, *I*(*H*_*t*_, *X*_*t*_). This prescription for encoding relevant collective variables can be formalized by the information bottleneck (IB) method (28, 30, 31). The information bottleneck objective function combines both of our stated goals – compression and predictive power – into one mathematical expression:

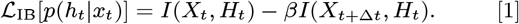

Crucially, the parameter *β* allows one to tune how much weight to assign to compression versus prediction. For small *β* the compression term dominates and the optimal encoder is trivial, losing all information about the system. For intermediate *β* the compression term does not allow *X*_*t*_ to be completely captured by *H*_*t*_, so that features of *X*_*t*_ must “compete” to pass through to the encoding variable (Fig. 1b). These features are reflected in the form of the encoder

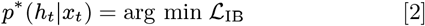

which provides the optimal trade-off between compression and predictability (29). Our goal is to connect the dynamical properties of the system to the features learned by the encoder.

In any realistic experimental setting, the presence of noise or uncertainty means we cannot predict precisely the future state of a system but instead can only predict a likely *distribution* of possible future states. Our prediction of the state at Δ*t* in the future is then represented mathematically as *p*(*x*_*t*+Δ*t*_|*x*_*t*_), the probability of observing state *x*_*t*+Δ*t*_ given the current state *x*_*t*_. This conditional probability distribution completely characterizes the dynamics of the system, and determines how probability distributions evolve in time:

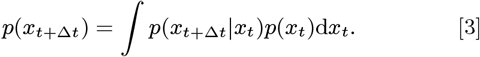

For Markovian, or “memoryless” dynamics, such an evolution can be understood as the action of a (linear) transfer operator *U* which acts on probability distributions. *U* can be decomposed into its right and left eigenvectors as

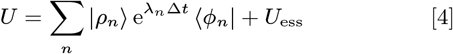

where |*ρ*_*n*_⟩ are right eigenvectors with eigenvalue 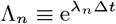 and ⟨*ϕ*_*n*_| are the corresponding left eigenvectors. *λ*_*n*_ are the eigenvalues of the infinitesimal generator of *U*, known as the Fokker-Planck operator (Fig. 1c). The operator *U*_ess_ corresponds to the so-called essential spectrum, and we assume that it can be neglected. This is usually possible when the system is subjected to even a small amount of noise, or when some amount of uncertainty is present in the measurements (33, 34). The eigenfunctions *ϕ*_*n*_ in Eq. (4) are in some sense “natural” features of the dynamics, as they evolve independently in time.

Our key observation is that the optimal encoder in Eq. (2) can be expressed in terms of the eigenvalues *λ*_*n*_ and left eigenfunctions *ϕ*_*n*_ of (the generator of) *U*,

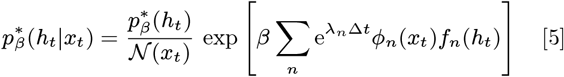

where *f*_*n*_(*h*_*t*_) are factors that do not depend on *x*_*t*_. For an outline of the mathematical steps leading to this see Methods, as well as the SI. These factors effectively determine what the encoder learns about the state *x*_*t*_. In general, there may be a large number of non-zero factors *f*_*n*_ so that the learned features are difficult to extract. However, things become simple in the limit of small *β*, or high compression. When *β* is small the encoder is trivial: *p*(*h*_*t*_|*x*_*t*_) = *p*(*h*_*t*_). In this case the value *h*_*t*_ is assigned at random with no regard to the state *x*_*t*_ of the system. No feature has been learned, and all factors *f*_*n*_ are equal to zero. As *β* is increased, the encoder undergoes a series of transitions at *β* = *β*_1_ < *β*_2_ < *β*_3_… where new features are allowed to pass through the bottleneck (Fig. 1b) (35–38). The first transition happens at a finite value of *β*_1_ when the first most predictive feature is learnt.

Surprisingly, we find that at the first IB transition the vector of *f*_*n*_ coefficients is dominated by a single term *f*_1_.

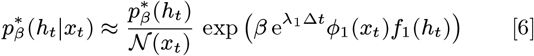

This is our main mathematical result, which we derive by considering a perturbative expansion of the IB objective for small *f*_*n*_. A proof of Eq. 6 with clearly specified technical assumptions may be found in the Methods and SI.

The above statement shows that in the limit of high compression the encoder’s dependence on *x*_*t*_ is given by the first left eigenfunction *ϕ*_1_(*x*_*t*_), which is the slowest-varying function of the state under dynamics given by *U*. Therefore, Eq. 6 makes precise the intuitive statement that slow features are the most relevant for predicting the future. Our analytical result, while applying only to the dominant eigenfunction, is valid for arbitrary non-Gaussian variables. The question of maximally informative features has additionally been explored in the context of animal vision, where one seeks to understand what features of the field of vision are encoded by retinal neurons (39, 40).

We further observe numerically that this picture holds true more generally: also at successive IB transitions, the learned features correspond to successive modes of the transfer operator. This picture is consistent with the exact results known for Gaussian IB, where the encoder learns eigenvectors of a matrix (related to the covariance of the joint *X*_*t*_, *X*_*t*+Δ*t*_ distribution) in a step-wise fashion at each IB transition (37). Together, this shows that the most informative features extracted by IB, an agnostic information-theoretic approach, correspond to physically-interpretable quantities – namely transfer operator eigenfunctions. As we show later, the insight above can be leveraged to systematically learn these slow variables directly from data with neural networks (41).

## 2. Information decay and the spectrum of the transfer operator

To develop intuition for information in a dynamical system, we turn to the simple example of a Brownian particle trapped in a confining double-well potential. This might represent, for example, a molecule with a single degree of freedom that transitions between two metastable configurations (42). In the overdamped limit the state of the particle is completely determined by its position *X*_*t*_ ∈ ℝ, with dynamics given by the Langevin equation

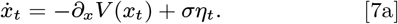

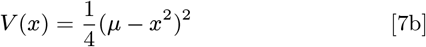

Here, *η*_*t*_ is unit-variance white noise, *σ* controls its strength, and *µ* controls the shape of the potential *V* (*x*).

The deterministic dynamical system undergoes a bifurcation at *µ* = 0 (Fig. 2a). Sample trajectories, with noise, for a uniform initial distribution of particles are shown in Fig. 2b. For negative *µ*, the trajectories all converge to a fixed point at *x* = 0, while for *µ* > 0 they fluctuate around the fixed points at 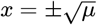.

**Fig. 2.**
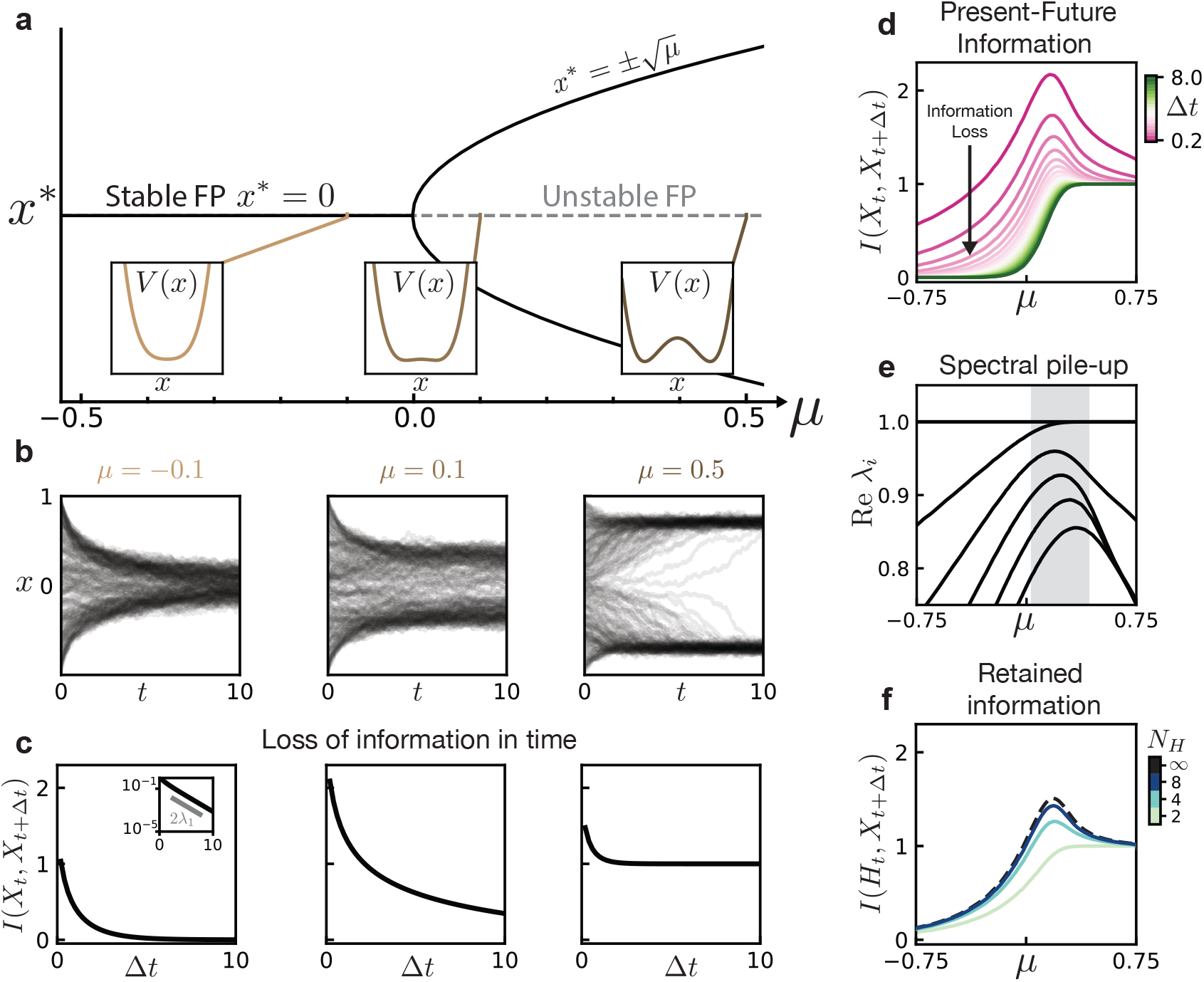
Information loss of a Brownian particle in a double well potential. (a) Fixed point (FP) diagram of the dynamics given by Eq. 7 for zero noise. There is a bifurcation at *µ* = 0 where the stable FP at *x* = 0 becomes unstable and two new stable FPs appear at 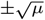. Insets show the evolution of the corresponding potential *V* (*x*), with the emergence of a double-well structure for *µ* > 0. (b) Dynamics of the system Eq. 7 for varying values of *µ* corresponding to the potential insets in (a), with uniformly-distributed initial conditions. (c) Loss of information between the initial condition and the future state. Inset shows scaling given by the first eigenvalue of the transfer operator. (d) Mutual information between the present and future state for varying time delay Δ*t* and bifurcation parameter *µ*. (e) Spectrum of the transfer operator *U*, showing a pile-up of eigenvalues for *µ* ≳ 0. These are related to the eigenvalues of its infinitesimal generator by 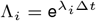 (f) Maximal mutual information which can be encoded into a discrete variable of *N*_*H*_ values, for a fixed time delay Δ*t* = 1.0. Black dashed line shows *I*(*X*_*t*_, *X*_*t*+Δ*t*_) for reference. Information is provided in units of bits.

To quantify the amount of information about the future state *X*_*t*+Δ*t*_ contained in the initial state *X*_*t*_ we compute their mutual information (Fig. 2c; see SI for details). The dynamics of *X*_*t*_ are Markovian, so that for any sequence of times 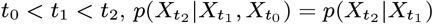. From the data processing inequality, one has (43)

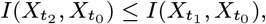

which implies that information can only decrease in time.

What governs the rate at which information decays? Here we can already see the role of the spectrum of the dynamics’ transfer operator. By exploiting the spectral expansion of the conditional distribution *p*(*x*_*t*+Δ*t*_|*x*_*t*_) one finds that f or long times the information decays as

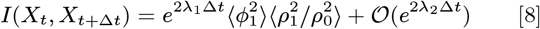

where expectations are taken over the steady state distribution (see SI). Asymptotically, the information decay is set by the value of *λ*_1_, the rate of decay of the slowest-varying function *ϕ*_1_(*x*) under the dynamics of *U*. In the limit of infinite time, for *any* value of *µ* even weak noise will cause the mutual information to become zero as there is a non-zero probability of hopping between the wells, though this rate of hopping is exponentially small (33).

The loss of information in time depends on the bifurcation parameter *µ* as summarized in Fig. 2d. Note the peak in *I*(*X*_*t*_, *X*_*t*+Δ*t*_) for small, positive *µ*. This corresponds to dynamics where observation of *X*_*t*_ strongly informs the future state; recall that the mutual information is maximized when the conditional entropy *S*(*X*_*t*+Δ*t*_|*X*_*t*_) ≈ 0 (see Methods). In contrast, for large positive or negative *µ, X*_*t*_ is not as informative of *X*_*t*+Δ*t*_ even for small times: the initial state is quickly forgotten as the particle approaches the bottom of the single (for *µ* < 0) or double (for *µ* > 0) well.

This phenomenon is reminiscent of critical slowing down, which occurs in the noise-free system as *µ* passes through the bifurcation at *µ* = 0. For the deterministic dynamics, the slowing down is reflected in the spectrum as a “pile up” of eigenvalues to form a continuous spectrum (33). In the presence of noise, although the continuous spectrum becomes discrete (33, 34) there is still a pile-up of eigenvalues characterized by several eigenvalues becoming close to 1 (Fig. 2e). This pile-up gives rise to the information peak seen in Fig. 2d. The peak is not solely due to the closing spectral gap *λ*_1_ − *λ*_2_, but is also impacted by the subdominant eigenvalues which accumulate at *µ* ≈ 0.2 (SI Fig. S4).

## 3. Knowing when to stop

For discrete encoding variables *h*, the information bottleneck partitions state space and reduces the dynamics on *x* to a discrete dynamics on *h*. Such reductions of complex systems to symbolic sequences via partitioning of state space has attracted attention for more than half a century in both theoretical and data-driven contexts(44–50). Several works have approached this partition problem from a dynamical systems perspective, linking “optimal” partitions to eigenfunctions of the (adjoint) transfer operator (25, 51). In this setting, a central question is “when to stop” (25, 26, 48, 49): how many states does *h* need in order to capture statistical properties of the original dynamics?

We consider this question by finding the optimal IB encoder in the limit of *low* compression, *β* ≫ 1, but fixed encoding capacity *N*_*H*_ (where *H*_*t*_ ∈ {0, …, *N*_*H*_ − 1}), i.e. the encoder is only restricted by the number of symbols it can use. An analogous setup was used in the context of renormalization group (RG) transformations in (52–54), which results in effective model reduction due to the “sloppiness”, or irrelevance, of certain system variables (55, 56). In this regime, the encoder learned by IB is deterministic; we are learning an optimal *hard* partition of state space. This can be seen by noting that *I*(*H*_*t*_; *X*_*t*+Δ*t*_) = *S*(*H*_*t*_) − *S*(*H*_*t*_|*X*_*t*+Δ*t*_) is maximized when the latter term is zero, which happens when *x*_*t*_ unambiguously determines *h*_*t*_, i.e. when *p*(*h*_*t*_|*x*_*t*_) ∈ {0, 1} for all *x*_*t*_. The details of how the encoder is computed are discussed in the next section.

Fig. 2f shows that the number of states necessary to describe the system depends strongly on the value of *µ*. For |*µ*| ≫ 0, a two-state discrete variable *h*_*t*_ ∈ {0, 1} suffices to describe the system’s future. Increasing the number of reduced variables *N*_*H*_ does not allow more information to be captured. Near the information peak at *µ* ≈ 0.2 this changes: predicting the future state of the system requires a more complex hidden variable of up to *N*_*H*_ ≈ 10 values. Above, we saw that this peak arises due to the pile-up of eigenvalues at *µ* ≈0.2. The content of the transfer operator spectrum is thus directly reflected in the number of encoding variables needed to capture the system’s statistics.

Noise can have a similarly dramatic impact on the reduced model complexity. Indeed, noise in some form, either inherent to the dynamics or due to measurement error, is necessary for a model to be reducible. In purely deterministic systems where the future state is a bijective function of the present state, information does not decay and complete knowledge of the state is required to predict the future.

Consider a fluctuating Brownian particle as above, where now each of the wells is split into two smaller wells, giving a total of four potential minima (Fig. 3a). As the system is in steady state, the standard deviation of the fluctuations *σ* corresponds to an energy scale *E*_*σ*_ = *σ*^2^ = 2*k*_B_*T*. For small *E*_*σ*_, the system rarely transitions between the four potential minima. In this case, knowledge of the initial minimum is very informative of the future state of the particle. In contrast, for large fluctuations the particle can spontaneously jump between shallow minima in each large well, so that the system immediately forgets about the precise potential minimum it was in. Information about the shallow minima has been “washed out”, and only the information about the larger double-well structure remains.

**Fig. 3.**
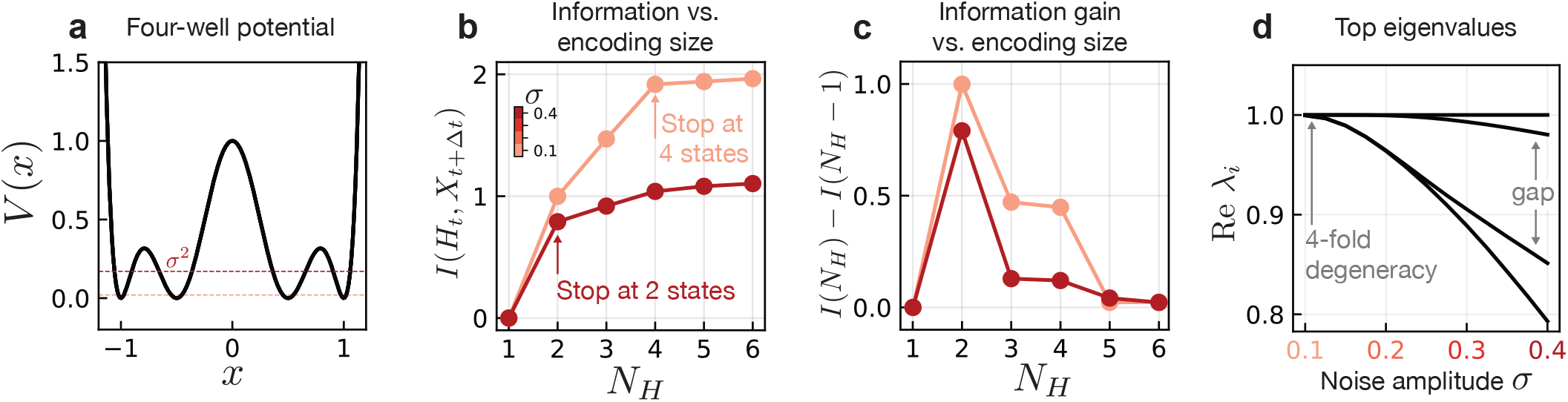
“Knowing when to stop”. The spectral properties of the transfer operator determine the necessary complexity (i.e. “when to stop” (25)) of the reduced model, which we show is also visible in information theoretic metrics. (a) Four-well potential in which a Brownian particle fluctuates. The magnitude *σ* of the fluctuating noise is related to an energy scale *E*_*σ*_ = *σ*^2^. (b) Information contained in the encoding variable *H*_*t*_ about the future state *X*_*t*+Δ*t*_ for varying levels of noise and alphabet sizes *N*_*H*_. (c) Information gain achieved by increasing the alphabet size by a single variable. This is the discrete derivative of the curve in (b). (d) Spectrum of the transfer operator for changing values of noise amplitude.

To see this reflected in the information, we again consider an encoding of the initial state into a discrete variable *H*_*t*_ ∈ {0, …, *N*_*H*_ − 1}. In both the small and large noise scenarios, a variable with *N*_*H*_ = 2 encodes approximately one bit of information (Fig. 3b), corresponding to an *H*_*t*_ which distinguishes the two large wells for *x* ≶ 0. For large noise this is essentially all the information that can be learned; increasing the capacity of the encoding variable beyond this provides only marginally more information about the future state (Fig. 3c). In the small noise case, the information between the encoding and the future state continues to increase to approximately two bits at *N*_*H*_ = 4, after which it plateaus. The encoding has learned to distinguish each of the four potential wells. These observations are reflected in the transfer operator spectrum shown in Fig. 3d. For small noise, the eigenvalue *λ* = 1 is four-fold degenerate which indicates the existence of four regions that can evolve independently under *U*, giving rise to four steady state distributions satisfying *Uρ* = *ρ*. These regions correspond to the potential minima. Hops between the separate minima are exceedingly rare, so that the dynamics essentially take place in the four minima independently. With larger *σ* the degeneracy is lifted, resulting in one dominant subleading eigenvalue followed by a gap. The corresponding eigenfunction is one which is positive (negative) on the right (left) side of the large potential barrier at *x* = 0: the only relevant piece of information is which of the large wells the initial condition is contained in, and all other information is lost exponentially quickly.

## 4. Transfer operator eigenfunctions are most informative features

Until now we have concerned ourselves with encodings whose capacity is limited only by the number of variables, rather than by the compression imposed by a small value of *β*. In the regime of small *β*, or high compression, features of the state *x*_*t*_ are forced to compete to make it through the bottleneck *h*_*t*_. By studying the behavior of the encoder in this regime, in particular its dependence on *x*_*t*_, we may identify the most relevant features of the state variable and show that they coincide with left eigenfunctions of the transfer operator.

We return to the simple example of a particle in a double well with dynamics given by Eq. (7) which we map to a discrete variable *H*_*t*_ ∈ {0, …, *N*_*H*_ − 1}. In this system the IB loss function Eq. (1) can be optimized directly, as shown in Ref. (30), using an iterative scheme known as the Blahut-Arimoto algorithm (43) (see SI).

To focus on the properties of encodings for varying degrees of compression *β*, we consider a fixed set of dynamical parameters *µ* and *σ*. Increasing *β* reduces the amount of compression, i.e. “widens” the bottleneck, allowing more information to pass into the encoder. This leads to a series of IB transitions which are sketched in Fig. 1b and shown quantitatively in Fig. 4a. The form of the optimal encoder changes qualitatively at these transitions. Before *β*_1_, the optimal encoder has no dependence on *x* so that *p*(*H*_*t*_ = *h*_*i*_|*x*_*t*_) = const for all *h*_*i*_. After the first transition, the encoder begins to associate regions of *x* to particular values of *h*. We are interested in the form of the encoder at *β* ≳ *β*_1_, just above the first IB transition, as this reflects the *most informative* features of the full state variable *x* (Fig. 4a). The dependence of *p*(*h*_*t*_|*x*_*t*_) on *x* can be explained by a stability analysis of the IB Lagrangian (see Methods and SI). Stability is governed by the eigenvalues *η*_*i*_ of the Hessian of the IB Lagrangian with respect to the parameters *f*_*n*_(*h*_*t*_) in Eq. 5. These parameters tell us how much the encoder “weights” each transfer operator eigenfunction; *f*_*n*_(*h*_*t*_) = 0 (for *n* > 0) corresponds to the uniform, or trivial encoder *p*(*h*_*t*_|*x*_*t*_) = *p*(*h*_*t*_).

**Fig. 4.**
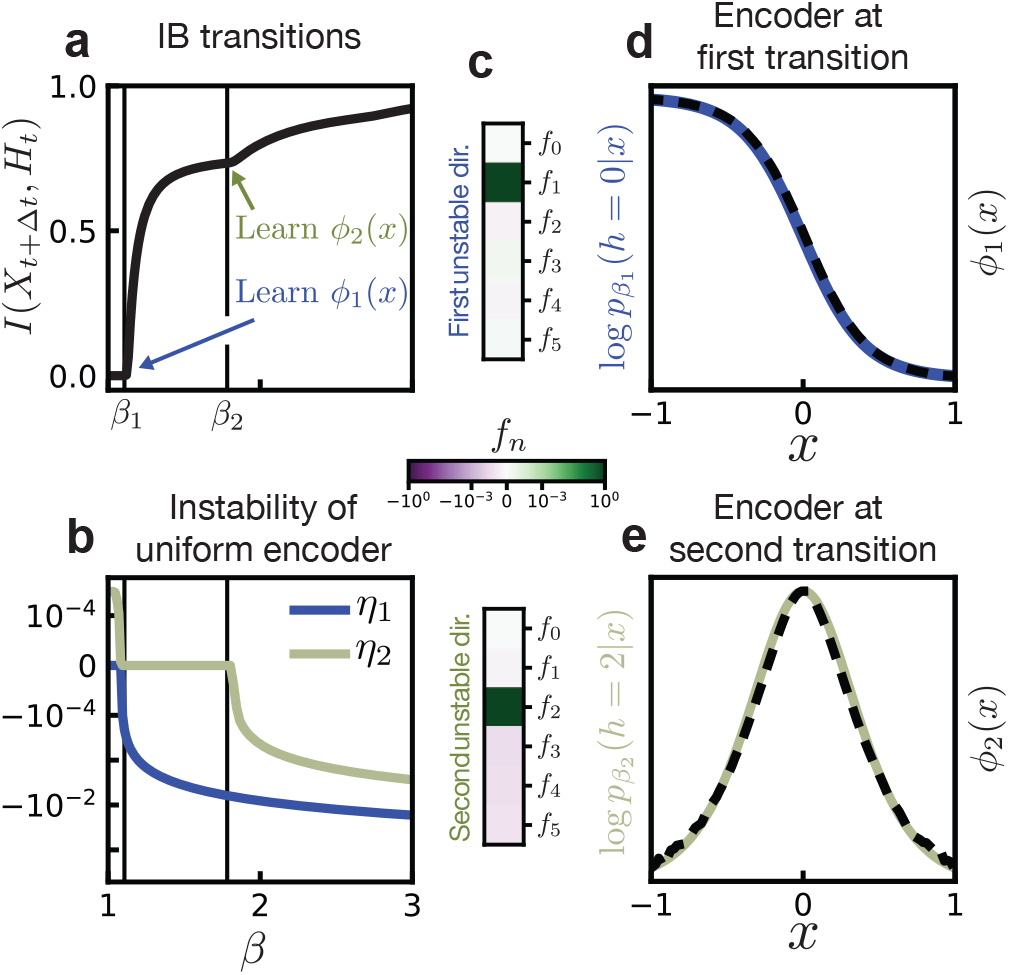
IB learns eigenfunctions of the adjoint transfer operator. (a) When the relative weight *β* between both constraints in Eq. 1 is changed, more and more information can go through the encoder. This occurs in steps, where the spectral content of the transfer operator is included starting from eigenvalues with largest magnitude (i.e., the slowest ones). (b) Transitions are characterized by the appearance of negative eigenvalues in the spectrum of the Hessian of the IB loss function. Here we consider the Hessian evaluated at the uniform encoder 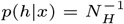. The IB transitions *β*_1_ ≈ 1.1 and *β*_2_ ≈ 1.8 correspond to the appearence of negative eigenvalues of the Hessian. (c) The unstable directions are dominated by single components (note the color scale is logarithmic). (d) At the first transition, the logarithm of the encoder is given by the eigenfunction *ϕ*_1_(*x*), up to rescaling (y-axis is shown in arbitrary units). (e) Likewise, at the second transition the encoder is given by *ϕ*_2_(*x*).

For small *β* all eigenvalues *η*_*i*_ are positive, indicating that the uniform encoder is a stable minimum of the IB Lagrangian. In Fig. 4b we show the smallest two eigenvalues of the IB Hessian when evaluated at the uniform encoder. At the first transition one eigenvalue becomes negative, so that the uniform encoder is unstable. The eigenvector corresponding to the unstable eigenvalue *η*_1_ indicates how the weights *f*_*n*_(*h*_*t*_) should be adjusted to lower the value of the IB Lagrangian. Our numerics confirm that these weights are dominated by *f*_1_ as expected from our analytical result (Fig. 4c, top). By taking the logarithm of the encoder after the transition, we can independently confirm that the encoder depends only on *ϕ*_1_(*x*) (Fig. 4d).

Our stability analysis predicts that a second mode becomes unstable at the second IB transition *β* ≈ *β*_2_ (Fig. 4b). Here we see that this unstable mode selects *f*_2_, and that the encoder correspondingly gains dependence on *ϕ*_2_(*x*) (Fig. 4d). Note that in general, *η*_2_ must not necessarily become negative precisely at *β*_2_ because the stability analysis is performed at the uniform encoder while the true optimal encoder has already deviated from uniformity. In the SI, we perform the same analysis for a triple-well potential where this difference is more apparent.

## 5. Data-driven discovery of slow variables

IB finds transfer operator eigenfunctions by optimizing an information theoretic-objective that makes no reference to physics or dynamics. This suggests it may be used for the discovery of slow variables in situations where one lacks physical intuition. The utility of exact IB for this purpose is limited because it requires knowledge of the exact conditional distribution *p*(*x*_*t*+Δ*t*_|*x*_*t*_) which is difficult to estimate in many real-world scenarios. Fortunately however, the IB optimization problem can be replaced by an approximate variational objective introduced in Ref. (41) that can be solved with neural networks. We refer to this as variational IB. In the remainder of this paper, we show how to implement these networks for the discovery of slow variables directly from data.

First we show numerically that the results of the previous sections remain valid for high-dimensional systems by considering a simulated data of fluid flow past a disk (57). The state of the system is given by a two-dimensional velocity field 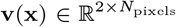, where *N*_pixels_ ∼𝒪 (10^5^) (Fig. 5a). Fluid flows in from the left boundary with a constant velocity *v*_0_*ê*_*x*_ past a disk of unit diameter. At Reynolds number Re ≳ 150, the fluid undergoes periodic vortex shedding behind the disk, forming what is known as a von Kármán street.

**Fig. 5.**
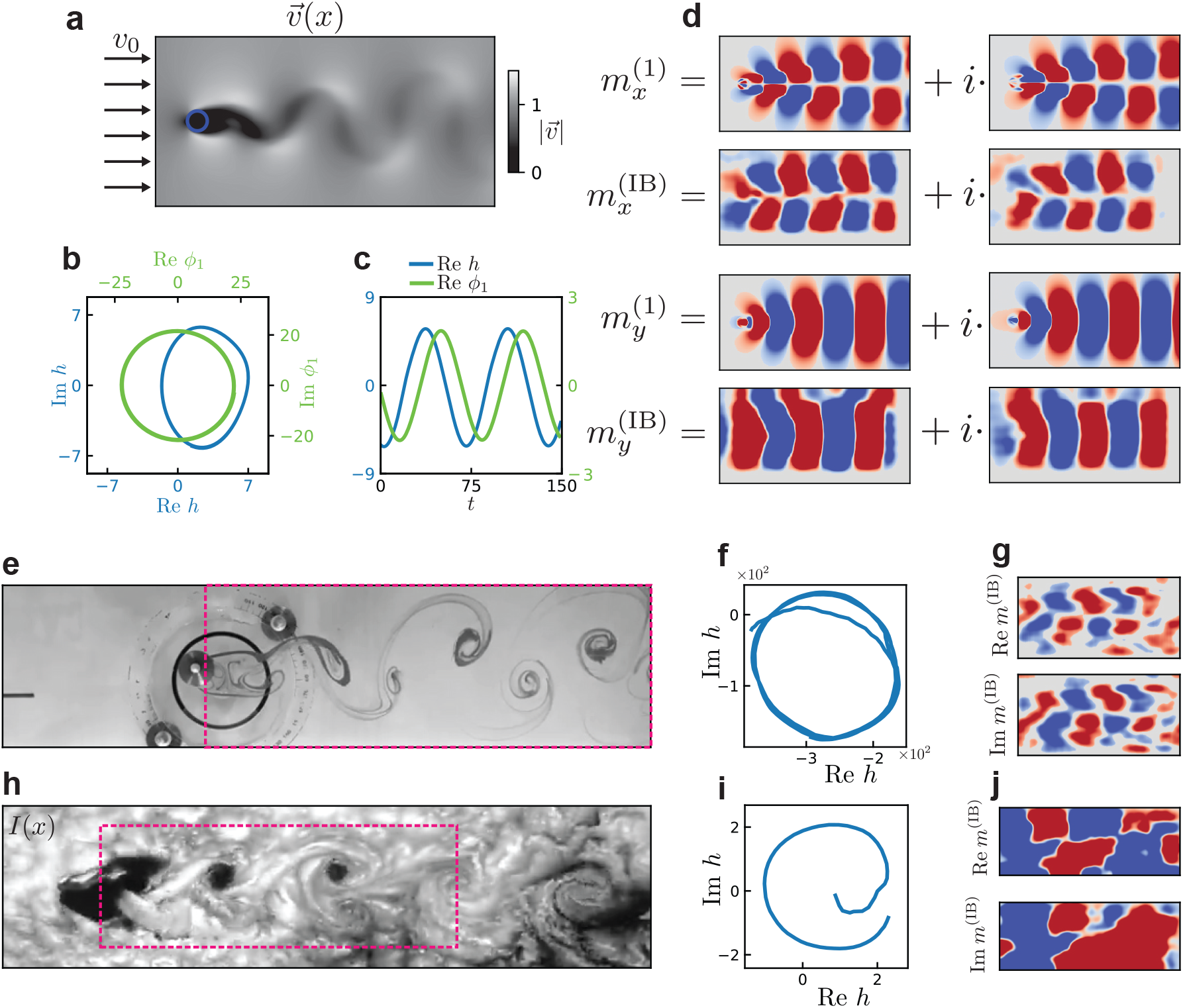
Variational IB for high-dimensional simulated fluid flow. (a) A fluid flows into the system with uniform velocity *v*_0_ in the *x*-direction and passes a disk-shaped obstacle, which perturbs the fluid and causes vortex shedding behind the object in a so-called von Kármán street. The state of the system is given by a spatially varying two-component vector field **v**(**x**). (b) The dynamics in latent space (blue) are very regular, traversing a nearly circular trajectory. For comparison we show the evolution of the mode amplitudes obtained by projecting the velocity field onto the first DMD mode (green). (c) Time evolution of one component of the latent variable (*h*_1_, blue) as well as the DMD mode amplitude (green). (d) Comparison of the first Koopman mode obtained from DMD (**m**^(1)^) and from VIB (**m**(^IB^)). Koopman modes from VIB are computed as gradients of the latent encoding variables as described in the main text. Red corresponds to positive values and blue to negative; the magnitudes of the modes are not directly comparable.

What do the true eigenfunctions look like in this system? Because it is well approximated by linear dynamics, eigen-functions of the adjoint transfer operator are linear functions of the state variable,

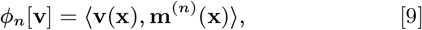

where **m**^(n)^ is the *n*-th mode (often referred to as a *Koopman* mode (19)) and angled brackets denote integration over space. The true eigenfunction and corresponding modes can be computed via dynamic mode decomposition (DMD) (11, 12), as described in the SI. The eigenfunctions for this system are in general complex, and come in conjugate pairs: 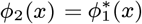. In this situation any linear combination of *ϕ*_1_ and *ϕ*_2_ will decay at the same rate, and hence we expect to learn some arbitrary combination of the two dominant eigenfunctions, or equivalently a combination of the real and imaginary parts of *ϕ*_1_. We therefore take a two-dimensional encoding variable [*h*_0_, *h*_1_], so that it can represent the full complex eigenfunction rather than only the real or imaginary part.

Our learned latent variables are oscillatory with the correct frequency as shown in Fig. 5b-c. A more stringent test is whether we are also learning the correct mode **m**^(1)^. From the true eigenfunctions, the modes can be extracted by computing the gradient

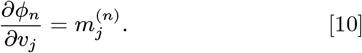

As the learned function *h*[**v**] is a neural network, we can efficiently compute gradients of the network with respect to the input field

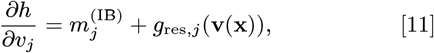

where we have separated the part of the gradient which is independent of **v** from a residual part which is dependent on **v**. If *h* corresponds to the true eigenfunction, we expect that **m**^(IB)^ is approximately equal to the Koopman mode **m**^(1)^, and that **g**_res_ is small. We indeed find this to be the case; Fig. 5d shows these gradients averaged over several instantiations of the neural network, which corresponds strongly to the true mode. Details concerning both the averaging procedure and the residuals **g**_res_ can be found in the SI. This shows that variational IB not only recovers the essential oscillatory nature of the dynamics, but does so by learning the correct slowly varying functions of the state variable given by the adjoint transfer operator eigenfunctions.

## 6. Relevant variable identification in laboratory-generated and atmospheric flows

The scenario above is characterized by high-dimensional data and few samples; training was performed with only ∼ 400 samples. We now demonstrate that our framework continues to hold approximately and yield interpretable latent spaces even for real-world fluid flow datasets scraped directly from videos on Youtube (58, 59) (Supplementary Movie 1).

The first shows a von Kármán street which forms as water passes by a cylindrical obstacle at Reynolds number 171, with flow visualized by a dye injected at the site of the obstacle (58). We take a background-subtracted grayscale image of the flow field as our input (Fig. 5e) and task VIB with learning a two-dimensional latent variable as above. Also here, variational IB learns oscillatory dynamics of the latent variables (Fig. 5f). We visualize the function learned by the encoder by considering gradients of the latent variables, which show the same structure as those obtained for the *x* component of the simulated data (Fig. 5g). This is expected, as the *x*-component of the velocity field has similar glide reflection symmetry as the intensity image.

Next, we apply variational IB to a von Kármán street arising due to flow around Guadalupe Island, which was imaged by a National Oceanic and Atmospheric Administration (NOAA) satellite (59) (Fig. 5h). The video consists of only 62 frames, and the von Kármán street undergoes a single oscillation. Even with this small amount of data, the variational IB neural network learns latent variables which capture this oscillation and have the expected dependence on the input variables (Fig. 5i-j). As in the first experimental example, the gradients of the encoding variables show the glide symmetry of *m*_*x*_ due to the symmetry of the intensity pattern in Fig. 5d. This symmetry is less clear in the component 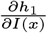, which is likely due to the fact that the von Kármán street is not as fully formed in this data as in our previous examples.

## 7. Relevant variable discovery in cyanobacterial populations

We now demonstrate how variational IB may be used as an aid for collective variable discovery in situations where physical intuition may not be a useful guide – collective behavior of biological organisms (Supplementary Movie 2). Here, we ask what the most predictive variables are for predicting the evolution of populations of cyanobacteria (S*ynechococcus elongatus*). The dynamics of the colonies are driven by several factors: growth and division of individual bacteria, translational motion of groups of bacteria as they are pushed by their neighbors, as well as the circadian oscillations within each bacterium (Fig. 6a). These oscillations are controlled by three Kai proteins (60) and depend in particular on the ratios of the copy number of these proteins which can be tuned experimentally (61).

**Fig. 6.**
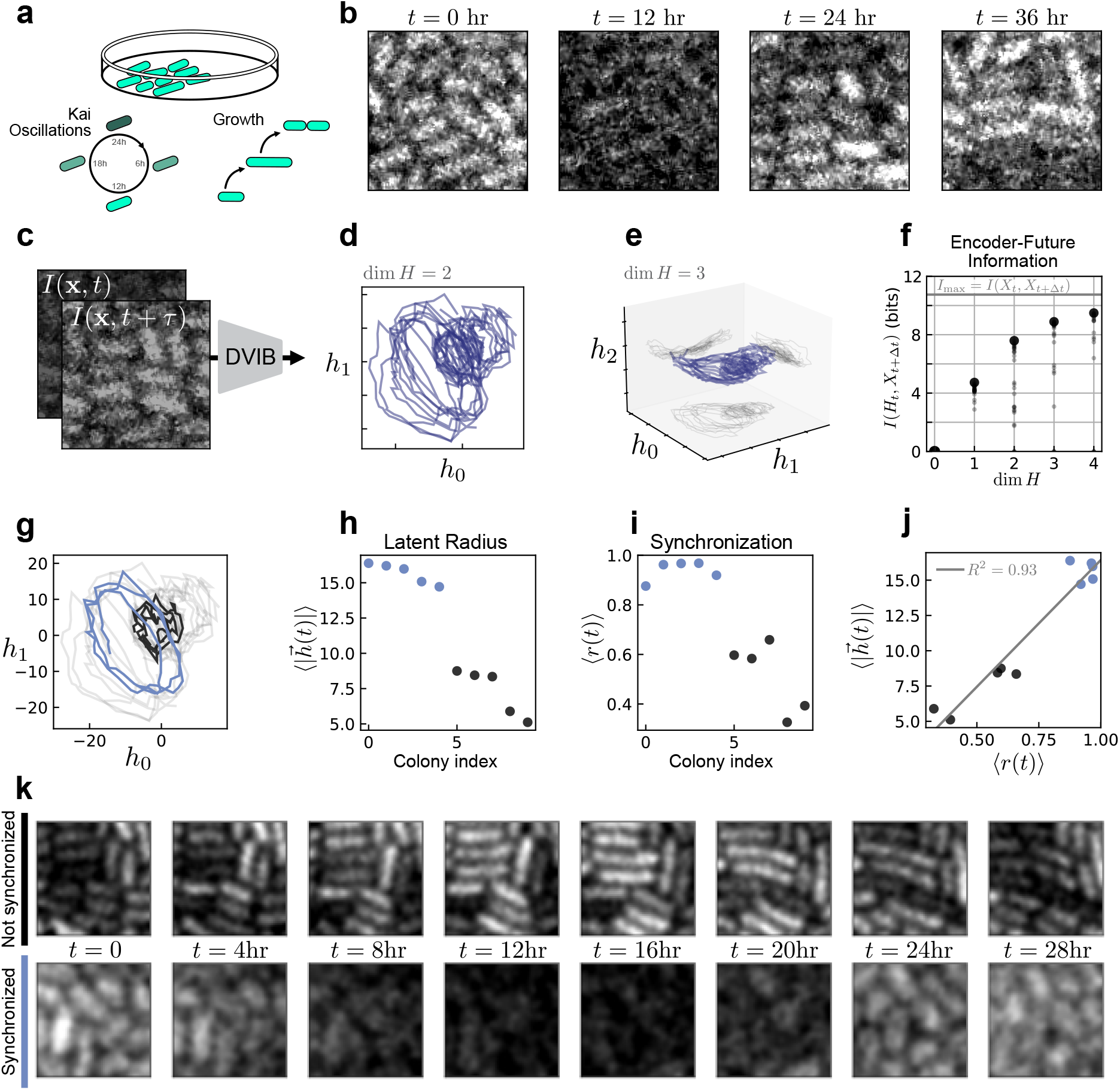
Discovering slow collective variables in cyanobacteria populations. (a) Fluorescent images of cyanobacteria colonies labeled with EYFP driven by the *kaiBC* promoter, allowing the visualization of Kai protein transcription. The colonies are imaged as they undergo cell growth and oscillations in Kai expression associated with the circadian rhythm. (b) Time series of one cropped section of an individual colony (see SI Fig. S9 for details). (c) The “state” used for variational IB is the time-lagged intensity field with lag time *τ*. (d) Variational IB embeddings of time-lagged images into two dimensions. Every line corresponds to one colony’s evolution in time. Note the apparent oscillations of different radii. (e) Embedding into three dimensions. We orient our axes to correspond to the three principal components of the data. Thus, the projection onto the *h*_2_ = 0 plane corresponds to a projection along the dominant two principal components. In this subspace, we see a similar structure as in the 2D embedding. (f) Mutual information between the future state and the encoding, given by *I*(*X*_*t*+Δ*t*_, *H*_*t*_), for varying dimension of the latent space. Increasing the dimension of the embedding space beyond two leads only to marginal increases in *I*(*X*_*t*+Δ*t*_, *H*_*t*_); this tells us “when to stop.” Each small point represents one training instance of the variational IB model, while the large point shows the maximum estimated value. Because the InfoNCE estimator is a lower bound on the true information, we consider the maximum as the estimated mutual information. The value of *I*_max_ = *I*(*X*_*t*_, *X*_*t*+Δ*t*_) is the mutual information estimated for the true dynamics. (g) Selected trajectories in latent space with large and small radii. (*h*) Time-average latent radius of all colonies. (i) Mean synchronization order parameter of the intensity images; see SI for computation details. (j) Mean radius versus synchronization parameter for each colony. VIB identifies clusters of cells characterized by high (blue) or low (black) synchronization. As revealed to us after our analysis, this clustering corresponds to differing theophylline concentrations across experiments. Within each experimental movie (each of which contains 2-3 colonies) the radii are mostly constant. (k) Sample time series of weakly (black) and highly (blue) synchronized colonies. We apply a slight Gaussian blur to better visualize the bacteria boundaries.

We were provided with videos of 10 cyanobacteria colonies that were grown under various conditions that impact their dynamics. However, as a test of our method, we were blinded to these conditions until we had performed our analysis. The videos are sequences of fluorescent images, taken once per hour, which show the clock state of each individual bacteria visualized with a fluorescent marker EYFP driven by the *kaiBC* promoter. Here, we focus on collective variables which are predictive of the state of the interior of the colony and not the growth in area of the colonies. We therefore crop the images to the interiors of each colony (SI Fig. S9). This allows us to isolate the motion of individual bacteria and fluorescence oscillations (Fig. 6b).

Our input to the variational IB neural network are these cropped images augmented with a time-lagged image of the same region (Fig. 6c). The purpose of this time lag is to make the dynamics Markovian: due to the oscillatory intensity field, if one observes only a single time point it is unclear whether the intensity is currently increasing or decreasing. These time lagged pairs comprise our system state, *X*_*t*_ ={*I*(**x**, *t*), *I*(**x**, *t* + *τ*)}, where *τ* is the duration of the time lag. Here we take *τ* = 3 hr and a prediction time horizon Δ*t* = 8 hr, but find that choosing different Δ*t* or *τ* does not change our results (SI Fig. S9).

With variational IB we compress the state *X*_*t*_ into a latent variable *h* of variable dimension (Fig. 6d-f). We train the neural network on the entire dataset of all 10 colonies simultaneously. The dynamics in latent space undergo clear oscillations, indicating that the relevant variables encode primarily the intensity fluctuations rather than, for example, the spatial locations of the bacteria. Notably, the trajectories are essentially two-dimensional, even when the encoding space is higher dimensional. This is reflected in the information retained about the future state, *I*(*X*_*t*+Δ*t*_, *H*_*t*_). We see that increasing the dimension of the embedding space beyond two leads only to marginal increases in *I*(*X*_*t*+Δ*t*_, *H*_*t*_); this tells us “when to stop” (Fig. 6f). We independently verify this by using principal component analysis to characterize the geometry of embedded trajectories, and find that even in higher dimensions the trajectories occupy a two dimensional subspace (SI Fig. S10). In the following, we therefore restrict our focus to the dim *H* = 2 case.

We noticed that there were notable differences in the radius of latent space oscillations from colony to colony, two of which are highlighted in (Fig. 6g). To understand this difference, we examined the original microscopy time series corresponding to both large and small latent radius (Fig. 6k) and found that while the large-radius sample showed clear, nearly uniform oscillations in intensity, the small-radius samples appeared much more heterogeneous.

To quantify this we consider each pixel to be an independent oscillator, akin to a spatial Kuramoto model (62–64), and compute a global synchronization order parameter *r*(t) (see SI). For each colony we calculate the time-averaged synchronization ⟨*r*(t)⟩_*t*_ and find that two clusters emerge corresponding to high and low synchronization (Fig. 6i). These clusters are precisely those representing trajectories of large and small latent radius (Fig. 6j), suggesting that variational IB learns to encode the synchronization of the colony in the latent variable radius. As a check, we perform IB on a simulated locally-coupled Kuramoto model as a system which shares many features of the experimental system. Here we also learn an encoding in which the latent radius corresponds to the synchronization order parameter (SI Fig. S11).

In the SI we compare the performance of variational IB to several other model reduction methods and find that IB delivers more interpretable and well-behaved features. This is likely due to the fact that many standard methods for data-driven model reduction rely on assumptions about the dynamics which may not be appropriate in the case at hand, such as linearity. Even among deep learning methods such as time-lagged autoencoders that are free of such assumptions, the variables learned by IB appear more interpretable. This increased interpretability is likely due to the compression term which effectively regularizes the latent space by encouraging the network to learn slow transfer operator eigenfunctions. While there are many specific use variants of DMD (13, 65–68) or autoencoders for dynamics (20–22, 69) that may outperform variational IB in some cases, we find that in this real-world example it yields the smoothest and most interpretable latent variables without requiring tailored pre-processing steps (SI Fig. S12).

By using variational IB we could reduce a complex system with multiple dynamical components – cell growth, division, and gene expression fluctuations – into a low dimensional form that retains only the most relevant information for the future. In addition to the insight that the dynamics are dominated by oscillations in two dimensions, the latent variables clearly distinguished trajectories into two groups that were not apparent *a priori*. We were provided this data as a “blind” test with no knowledge of the underlying system. After we performed our analysis, it was revealed to us that these bacterial colonies have been engineered to control the translational efficiency of the Kai proteins by varying theophylline concentration (61). The synchronization order parameter discovered by variational IB corresponds to differing experimental concentrations of theophylline, which is in agreement with the findings in Ref. (61). IB can thus serve as a way to connect experimental control parameters to effective changes in dynamics.

## 8. Conclusion

We have related information-theoretical properties of dynamical systems to the spectrum of the transfer operator. We illustrate our findings on several simple and analytically tractable systems, and turn them into a practical tool using variational IB, which learns an encoding variable with a neural network. The latent variables of these networks can be interpreted as transfer operator eigenfunctions even though the network was not explicitly constructed to learn these: it optimizes a purely information-theoretic objective that contains no knowledge of a transfer operator or dynamics. This allows one to harness the power of neural networks to learn physically-relevant latent variables. Biological systems are an ideal setting for such methods: despite their apparent complexity, they can often be captured by low-dimensional descriptions which are difficult to identify by physical considerations alone (47, 70–72). We have shown that variational IB is a potentially powerful tool for these cases, and can discover slow variables even directly from image data without significant preprocessing.

## Materials and Methods

### Mutual information and entropy

Let *X* be a random variable which takes values *x* that are observed with probability *p*(*x*). The entropy of this distribution measures the predictability of the outcome of a measurement of *X* and is given by (43)

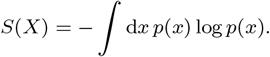

Given another random variable *Y*, such that *X* and *Y* have a joint distribution *p*(*x, y*), we can ask how much information is shared between these two variables. This is given by the mutual information

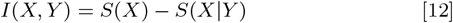

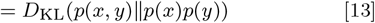

This can be interpreted as quantifying how much (on average) a measurement of *Y* can reduce our uncertainty about the value of *X* (Eq. (12)).

### The information bottleneck

The information bottleneck (30) is an example of a rate-distortion problem which seeks to find an optimal compression which minimizes some distortion measure with the original signal (43). Concretely, we call *X* the source signal, and let *H* denote the compressed signal. In IB, rather than using an *a priori* unknown distortion function, one seeks to ensure that the compression retains information about an additional relevance variable *Y*. As noted in the main text, the IB optimization objective is given by the Lagrangian

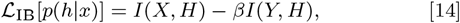

where in our case the source signal *X* is the state of the system *X*_*t*_ at time *t*, and the relevance variable is the state of the system *X*_*t*+Δ*t*_ at a future time *t* + Δ*t*. The encoder which optimizes this objective can be solved for exactly and is given by (30)

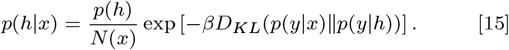

### Encoder in terms of transfer operator eigenfunctions

To connect the optimal encoder to the transfer operator, we first rewrite Eq. (15) in terms of the transition probabilities,

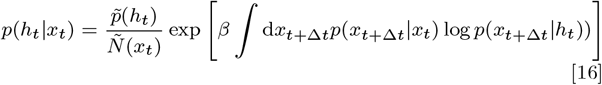

where we have absorbed terms in the exponent which depend only on *h*_*t*_ or *x*_*t*_ into the normalization factors. Into the above equation, we replace the transition probability with the spectral decomposition

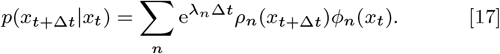

From this, the Eq. (5) of the main text immediately follows, where

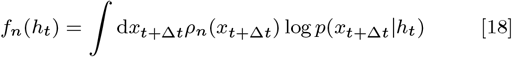

which may be interpreted as a sort of cross entropy (*ρ*_*n*_ is generally not a probability distribution) between each right eigenfunction and the decoding of *h*_*t*_ into the future state *x*_*t*+Δ*t*_.

To study the behavior of the encoder in the limit of high compression, we consider a transfer operator *U* with infinitesimal generator ℒ_*U*_. For ℒ_*U*_ with a discrete spectrum with eigenvalues satisfying 0 = *λ*_0_ > *λ*_1_ > *λ*_2_ ≫ *λ*_3_… and for *β* just above the first IB transition *β*_1_, we show that the optimal encoder is given approximately by

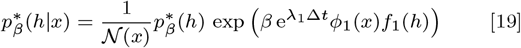

with corrections due to the second eigenfunction given by *f*_2_(*h*) ≈ *f*_1_(*h*)e^*−*ΓΔ*t*^ + 𝒪(e^*−*2ΓΔ*t*^) where Γ = *λ*_1_ *− λ*_2_ > 0 denotes the spectral gap. To see this, we compute the Hessian of the IB Lagrangian

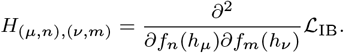

Here we assume a finite alphabet of size *N*_*H*_, i.e. *h*_*µ*_ with *µ* ∈ {1, …, *N*_*H*_}. At the uniform encoder, i.e. *f*_*n*_(*h*) = 0, the Hessian decomposes into a tensor product

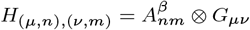

where *A*^*β*^ depends only on the indices of the coefficients and *G* captures the dependence on *h*_*ν*_. The only part which depends on *β* is *A*^*β*^.

We are concerned with the sign of the eigenvalues of *H*. A negative eigenvalue indicates that ℒ_IB_ is unstable to a perturbation in *f*, which means the loss can be lowered by changing *f* away from the trivial encoder at *f*_*n*_ = 0. Because the eigenvalues of a tensor product of matrices are products of the eigenvalues of the component matrices, the eigenvalues of *H* change sign when those of *A*^*β*^ do. *A*^*β*^ and its spectrum can be computed, which we do in the SI. The result of this calculation is that the first eigenvalue to become negative is associated with the eigenvector **v** = (1, 0, 0, …). This computation is exact for equilibrium systems, which are those in which the steady-state flux vanishes, but in nonequilibrium systems there may generally be a correction proportional to the flux. In summary, this means that in the limit of high-compression only the first component *f*_1_ becomes non-zero, hence the encoder has the form given in Eq. (19).

### Variational IB compared to other dimensionality reduction techniques

Variational IB (VIB) is by no means the only numerical method for performing data-driven model reduction. Here we provide a brief overview of the benefits and shortcomings of VIB with respect to other methods; an extended discussion can be found in the SI.

One class of methods is based on linear projections, such as principal component analysis (PCA), dynamic mode decomposition (DMD) (11, 12), or (time-lagged) independent component analysis (TICA) (10) (which is equivalent to DMD (73)). These methods can be extended to take into account non-linearity by introducing a library of non-linear terms on which one then applies the above methods, such as in kernel PCA (74) or extended DMD (eDMD) (13). These methods have the advantages, relative to VIB, that their optimization (even for the extended algorithms) relies only on linear projections which are fast and interpretable. However, the success of these methods depends on the choice of an appropriate library of functions so that the projection onto this space is closed under the dynamics. Choosing an appropriate library is not always possible (75, 76).

A second category of non-linear dimensionality reduction techniques are graph-based or similarity-based methods, which typically assume that the data is distributed on a low-dimensional manifold embedded in a higher-dimensional space (77, 78). One prominent example is diffusion maps (14), which starts from a set of data snapshots and, assuming the system evolves diffusively on short times, constructs an approximate transition matrix from which one can compute eigenfunctions to parameterize the data manifold. The assumption of diffusive dynamics can be violated when data is not sampled sufficiently frequently. This likely explains our finding that VIB produced more well-behaved low-dimensional embeddings on the cyanobacteria dataset (SI Fig. S12). VIB has the additional advantage, relative to this and similar methods, that it explicitly takes dynamics into account without the strong assumptions required by diffusion maps.

Finally, deep neural networks can be used for model reduction through encoder-decoder architectures that attempt to reconstruct the data from a low-dimensional latent space; VIB falls into this class of methods. Some standard neural network architectures from this class include autoencoders (AEs) and variational autoencoders (VAEs). For dynamical systems in particular, extensions to these methods have been proposed which impose constraints on the latent dynamics, such as linearity (20–22, 69). Autoencoders often produce poorly-behaved latent spaces that distribute the latent variables on a narrow manifold with sharp features, see for example (69). By regularizing the latent embedding to encourage smoothness, variational autoencoders can remedy some of these issues. We note that the VIB loss is very similar to a VAE loss with the contrastive InfoNCE loss replacing the reconstruction loss, so we expect that for many problems these should perform similarly. Other dynamically-constrained architectures such as in (20–22) work well for deterministic systems but it is unclear what effect stochasticity has on their performance. In our examples we have seen that VIB works well on noisy data.

In general when investigating a new system it is good practice to start by attempting to perform dimensionality reduction with linear methods such as PCA or DMD because they are fast, straightforward to implement, and easy to interpret. In situations where linear techniques are not sufficient, VIB may be preferable to other methods because it is guaranteed to find dynamically relevant variables (in contrast to diffusion maps, t-SNE, AEs, VAEs, etc.) and it does not require that one performs the carefully tailored preprocessing steps that are required by eDMD or kernel PCA, or other variants of DMD (65–68). Additionally, it works well even when the dynamics are highly stochastic as shown in the cyanobacteria dataset.

## ACKNOWLEDGMENTS

The authors would like to thank Z. Ringel, M. Han and D.E. Gökmen for helpful discussions. M.S.S. acknowledges support from a MRSEC-funded Graduate Research Fellowship (DMR-2011854). M.K.-J. gratefully acknowledges financial support from the European Union’s Horizon 2020 programme under Marie Sklodowska-Curie Grant Agreement No. 896004 (COMPLEX ML). D.S.S. acknowledges support from a MRSEC-funded Kadanoff–Rice fellowship and the University of Chicago Materials Research Science and Engineering Center (DMR-2011854). V.V. acknowledges support from the Simons Foundation, the Complex Dynamics and Systems Program of the Army Research Office under grant W911NF19-1-0268, the National Science Foundation under grant DMR-2118415 and the University of Chicago Materials Research Science and Engineering Center, which is funded by the National Science Foundation under award no. DMR-2011854. All authors acknowledge support from the UChicago Research Computing Center which provided the computing resources for this work.

