## Supplementary Information for "Information theory for data-driven model reduction in physics and biology"

#### This PDF file includes:

Supporting text
Figs. S1 to S13
SI References

### Supporting Information Text

### 1. Information bottleneck

The information bottleneck was originally formulated in Ref. (1) as a rate distortion problem. Rate distortion theory describes how to maximally compress a signal (i.e. minimize its communication rate) such that it remains minimally distorted, which, however, presupposes the knowledge of a distortion function specifying which features need to be preserved(2). In Ref. (1), Tishby et al. introduce a variant of the problem where the a priori unknown distortion function is not used, but rather one seeks to ensure that the compression retains information about an auxiliary variable $Y$ correlated with the signal. As the correlations with $Y$ define implicitly the relevant features of the signal to be preserved, $Y$ is called the relevance variable. Concretely, we call $X$ the source signal, and $H$ denotes the compressed signal. The random variables form a Markov chain $H \leftrightarrow X \leftrightarrow Y$, meaning that $H$ and $Y$ are conditionally independent given $X$:

$$p(y|h,x) = p(y|x)p(h,x)$$
$$p(h|x,y) = p(h|x)p(y,x)$$

As noted in the main text, the IB optimization objective is given by the Lagrangian

$$\mathcal{L}_{\rm IB}[p(h|x)] = I(X,H) - \beta I(Y,H). \quad [1]$$

To enforce normalization of the $p(h|x)$ one introduces a Lagrange multiplier $\lambda(x)$ so that the full optimization function is

$$\mathcal{L}_{\rm IB}[p(h|x)] = \sum_{x,h} p(h|x)p(x)\log\frac{p(h|x)}{p(h)} - \beta\sum_{y,h} p(y,h)\log\frac{p(y,h)}{p(h)p(y)} + \sum_x \lambda(x)\left(1-\sum_h p(h|x)\right). \quad [2]$$

The encoder which optimizes this objective can be solved for exactly. Using the following functional derivatives,

$$\frac{\delta}{\delta p(h|x)}p(h') = \delta(h-h')p(x)$$
$$\frac{\delta}{\delta p(h|x)}p(h',y') = \delta(h-h')p(x,y'),$$

one can compute the derivative of the Lagrangian. By evaluating the derivative and setting to zero, one finds

$$\log p(h|x) = \log p(h) - \beta\sum_y p(y|x)\log\frac{1}{p(y|h)} - \lambda(x)$$

which can be rearranged to give the optimal encoder

$$p(h|x) = \frac{p(h)}{N(x)}\exp\left[-\beta D_{KL}(p(y|x)\|p(y|h))\right]. \quad [3]$$

By absorbing terms which only depend on $h$ and $x$ into $p(h)$ and $N(x)$, respectively, the encoder can be expressed as

$$p(h|x) = \frac{p(h)}{N(x)}\exp\left[\beta\int dy\, p(y|x)\log p(y|h)\right]. \quad [4]$$

When $\beta < 1$, it follows from the data processing inequality $I(X,H) \geq I(Y,H)$ that Eq. (1) is minimized by a trivial encoder $p(h|x) = p(h)$. In this case, $\mathcal{L}_{\rm IB} = 0$ and no information passes through the bottleneck. As $\beta$ is increased, more information is allowed through the bottleneck until the relevant variables begin to gain a dependence on the state $x$. This occurs suddenly for a certain value of $\beta = \beta_1 > 1$ at which the encoder becomes non-uniform and $I(H,X)$ becomes non-zero. This is referred to as an IB transition; for increasing $\beta$, there may be a sequence of transitions at $\beta_2, \beta_3, \ldots$ etc.

### 2. Numerically solving "exact" IB and Ulam approximations of the transfer operator

The optimal IB encoder Eq. (3) can be found using the Blahut-Arimoto (BA) algorithm (2). As described in detail in Refs. (1, 3) and sketched in Fig. S1, the algorithm is an iterative procedure, where the encoder at iteration $k+1$ is updated according to

$$\begin{cases} p_{k+1}(h|x_t) = \frac{p_k(h)}{N_{k+1}(x)}\exp(-\beta D_{\rm KL}(p(x_{t_\Delta t}|x_t)\|p_k(x_{t+\Delta t}|h))) \\ p_{k+1}(h) = \sum_{x_t} p(x_t)p_{k+1}(h|x_t) \\ p_{k+1}(x_{t+\Delta t}|h) = \sum_{x_t} p(x_{t+\Delta t}|x_t)p_{k+1}(x_t|h). \end{cases}$$

The first line simply plugs the previous estimate of the encoder in to Eq. 10 (and normalizes the distribution), while the following two lines update marginal and conditional distributions using the new estimate of the encoder. In Refs. (1, 3) it is shown that this algorithm converges.

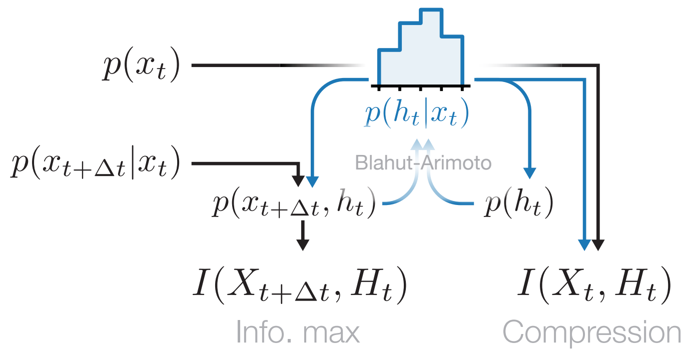


**Fig. S1. Exact IB.** Exact IB finds an optimal encoder using an iterative Blahut-Arimoto algorithm, described in the main text, for a rate-distortion problem with Kullback-Leibler distortion (1). This process requires access to (an estimate of) the transfer matrix and the steady state distribution. Once an optimal encoder has been found, relevant quantities such as mutual information can be computed.

The BA algorithm requires access to the conditional distribution $p(x_{t+\Delta t}|x_t)$ for each $x_t$, as well as the steady state $p(x_t)$. To solve the IB optimization problem we therefore need a numerical approximation of the transfer operator, which we obtain by an Ulam approximation (4, 5). In brief, one divides space into bins and computes a finite-dimensional approximation to the conditional distribution as

$$P_{ij} = P(X_{t+\Delta t} = x_j | X_t = x_i) = N_{i\to j}/N_i, \quad [5]$$

where $N_i$ is the number of trajectories starting in bin $i$ and $N_{i\to j}$ the number of observed transitions from bin $i$ to bin $j$. The transfer operator is then approximately given by

$$Up(x_j) \approx \sum_{x_i} P_{ij} p(x_i), \quad [6]$$

and eigenvalues and eigenvectors of $U$ can be computed by diagonalizing $P$.

### 3. Transfer operators

In this work we consider dynamical systems given by a Langevin equation

$$\dot{\vec{x}} = \vec{f}(\vec{x}) + \sqrt{2D(\vec{x})}\vec{\xi}(t) \quad [7]$$

where the first term is the deterministic part of the dynamics, and the second term corresponds to noise, where

$$\langle \xi_i(t)\xi_j(t')\rangle = C_{ij}\delta(t-t').$$

This framework captures purely deterministic dynamics, which are obtained by setting $D = 0$. The transfer operator describes the evolution of probability distributions. For dynamics given by a Langevin equation, probability distributions evolve according to the corresponding Fokker-Planck equation

$$[\mathcal{L}\rho](\vec{x}) = \partial_t \rho(\vec{x}) = -\partial_i(f_i(\vec{x})\rho(\vec{x})) + \partial_i\partial_j(D(\vec{x})C_{ij}\rho(\vec{x})).$$

Here we recognize $\mathcal{L}$ as the infinitesimal generator of the transfer operator $U$: evolution for a time $t$ is given by $U = \mathrm{e}^{t\mathcal{L}}$.

The adjoint operator is given by the so-called backward Kolmogorov equation, and it describes the evolution of functions $\phi$.

$$\mathcal{L}^\dagger \phi(x) = f_i(\vec{x})\partial_i(\phi(\vec{x})) + D(\vec{x})C_{ij}\partial_i\partial_j(\phi(\vec{x})). \quad [8]$$

For $D = 0$, $\mathcal{L}$ is the generator of the Perron-Frobenius operator, while its adjoint is the Koopman operator (6).

### 4. Derivations

Here we derive the form of the optimal encoder at the first IB transition,

$$p^*_\beta(h|x) \approx \frac{1}{\mathcal{N}(x)} p^*_\beta(h) \exp\left(\beta\, \mathrm{e}^{\lambda_1 \Delta t}\phi_1(x) f_1(h)\right) \quad [9]$$

In Ref. (1), it is shown that the encoding minimizing the IB objective satisfies the implicit equation

$$p^*_\beta(h|x) = \frac{p^*_\beta(h)}{\mathcal{N}(x)} \exp\left(-\beta D_{\mathrm{KL}}\left[p_{X_{t+\Delta t}|X_t} \middle|\middle| p_{X_{t+\Delta t}|H_t}\right]\right) \quad [10]$$

where $\mathcal{N}$ is a normalization factor ensuring $\int \mathrm{d}h\, p^*_\beta(h|x) = 1$, and

$$p^*_\beta(h) \equiv p^*_\beta(h) = \int \mathrm{d}x\, p^*_\beta(h|x) p(x). \quad [11]$$

The Kullback-Leibler divergence between two probability distributions $p(x)$ and $q(x)$ is defined by

$$D_{\mathrm{KL}}[p||q] = \int \mathrm{d}x\, p(x)\, \log \frac{p(x)}{q(x)}; \quad [12]$$

in the $D_{\mathrm{KL}}$ in Eq. (10), the integration is done with respect to the common random variable $X_{t+\Delta t}$ of the two conditional distributions.

Equation 10 contains the conditional distribution $p_{X_{t+\Delta t}|H_t}$, which is the composition of two operations: decoding (going from $H_t$ to $X_t$) and time evolution (going from $X_t$ to $X_{t+\Delta t}$). The decoding is performed using Bayes' theorem $p_{X_t|H_t} p_{H_t} = p_{H_t|X_t} p_{X_t}$, so that we have

$$p_{X_{t+\Delta t}|H_t} = \int_{X_t} p_{X_{t+\Delta t}|X_t}\, p_{H_t|X_t}\, \frac{p_{X_t}}{p_{H_t}}. \quad [13]$$

Note the appearance of the encoder $p_{H_t|X_t}$, making Eq. (10) only an implicit solution of the optimization problem.

The conditional distribution $p_{X_{t+\Delta t}|X_t}$ can be written in terms of right and left eigenvectors of $U$. Here we neglect $U_{\mathrm{ess}}$, which is the operator corresponding to the essential spectrum. The essential spectrum is the part of the spectrum that is not the discrete spectrum; by neglecting it, we are assuming that the essential radius $\rho_{\mathrm{ess}}$ (the maximum absolute value of eigenvalues in the essential spectrum) is small enough compared to the first few eigenvalues $|\lambda_n|$ in the point spectrum. While purely deterministic systems may exhibit a large essential radius, the introduction of noise causes the essential spectrum to shrink or disappear (7, 8). We can then write the conditional distribution in terms of right and left eigenvectors of $U$,

$$p_{X_{t+\Delta t}|X_t}(x|x') = \sum_n \mathrm{e}^{\lambda_n \Delta t} \rho_n(x) \phi_n(x'). \quad [14]$$

The Kullback-Leibler divergence in Eq. (10) can be represented as a sum of two terms,

$$D_{\mathrm{KL}}[...] = \int \mathrm{d}x_{t+\Delta t}\, p(x_{t+\Delta t}|x_t)\, \log p(x_{t+\Delta t}|x_t) - \int \mathrm{d}x_{t+\Delta t}\, p(x_{t+\Delta t}|x_t)\, \log p(x_{t+\Delta t}|h_t). \quad [15]$$

The first term has no dependence on $h$ and can hence be absorbed into the normalization $\mathcal{N}(x)$. Plugging the decomposition Eq. (14) into the second term leads directly to

$$p^*_\beta(h|x) = \frac{1}{\mathcal{N}(x)}\, p^*_\beta(h)\, \exp\left[\beta \sum_n \mathrm{e}^{\lambda_n \Delta t} \phi_n(x) f_n(h)\right] \quad [16]$$

where

$$f_n(h) = \int \mathrm{d}x_{t+\Delta t}\, \rho_n(x_{t+\Delta t}) \log p(x_{t+\Delta t}|h), \quad [17]$$

which can be understood as a quasi-cross entropy between the right eigenvector $\rho_n$ and the distribution on $x_{t+\Delta t}$ obtained by decoding $h$. Equation 16 depends only on the existence of the spectral decomposition; we have made no other assumptions on the dynamics (beyond Markovianity) up until this point.

**Perturbation theory at the uniform encoder.** In this section we derive our main result for the optimal encoder in the limit of high compression (small $\beta$).

*Mathematical result – Consider a transfer operator $U$ with infinitesimal generator $\mathcal{L}_U$. Assume that $\mathcal{L}_U$ has a discrete spectrum with eigenvalues satisfying $0 = Re\ \lambda_0 > Re\ \lambda_1 > Re\ \lambda_2 \gg Re\ \lambda_3 \ldots$. Then, for $\beta$ just above the first IB transition $\beta_1$ so that $f_n(h) \to 0$, the optimal encoder is given approximately by*

$$p^*_\beta(h|x) = \frac{1}{\mathcal{N}(x)} p^*_\beta(h)\, \exp\left(\beta\, \mathrm{e}^{\lambda_1 \Delta t} \phi_1(x) f_1(h)\right) \quad [18]$$

*with corrections due to the second eigenfunction given by $f_2(h) \approx f_1(h) e^{-\Gamma \Delta t} + \mathcal{O}(e^{-2\Gamma\Delta t})$ where $\Gamma = \lambda_1 - \lambda_2 > 0$ denotes the spectral gap.*

To show this we compute the $f_n(h)$ coefficients at the onset of instability. The instability at the first IB transition at $\beta = \beta_1$ corresponds to the emergence of negative eigenvalues in the Hessian of the IB loss,

$$\frac{\partial^2}{\partial f_n(h_i) \partial f_m(h_j)} \mathcal{L}_{\mathrm{IB}}. \quad [19]$$

Similar perturbative approaches have been studied in other contexts within IB in (9–11). For concreteness, we consider a discrete state $x \in \{1, ..., N_X\}$ and discrete encoding $h \in \{1, ..., N_H\}$. We express the encoder as in Eq. 16 where we now expand the normalization terms

$$p^*_\beta(h_\mu|x_i) = \frac{f_0(h_\mu)\exp\left[\beta \sum_n e^{\lambda_n \Delta t}\phi_n(x_i) f_n(h_\mu)\right]}{\sum_{h_\lambda} f_0(h_\lambda)\exp\left[\beta \sum_n e^{\lambda_n \Delta t}\phi_n(x_i) f_n(h_\lambda)\right]} \tag{20}$$

In the following, we simplify notation as

$$p(h_\mu|x_i) = \frac{f_0^\mu \exp\left[\sum_n \tilde{\phi}_n(x_i) f_n^\mu\right]}{\sum_{h_\lambda} f_0^\lambda \exp\left[\sum_n \tilde{\phi}_n(x_i) f_n^\lambda\right]} \tag{21}$$

where $\tilde{\phi}_n(x_i) = \beta e^{\lambda_n \Delta t}\phi_n(x_i)$. For the uniform encoder, $f_n^\mu = 0$ for all $n \neq 0$.

Hereafter we assign $\tilde{\phi}_0 = \frac{1}{f_0^\mu}$, taking care to track the correct Greek index which otherwise is not present for $\tilde{\phi}_n$ (in the following, $n$ is always paired with $\mu$ and $m$ always with $\nu$). For example, a single derivative of the marginal is given by

$$\partial_n^\mu p(h_\nu) = \sum_{x_i} p(x_i) p(h_\nu|x_i)\tilde{\phi}_n(x_i)\left(\delta_{\mu\nu} - p(h_\mu|x_i)\right). \tag{22}$$

To compute the Hessian, we use the fact that $\sum_\lambda \partial_n^\mu p(h_\lambda) = 0$ (by exchanging sum and derivative), and that $p(h_\mu|x_i)|_{\vec{f}=0} = p(h_\mu)$. We note in passing that we find the first derivative terms of $\mathcal{L}_{\mathrm{IB}}$ vanish, see also (11, 12). The compression term is given by

$$\begin{aligned}\partial_n^\mu \partial_m^\nu I(X,H) = &\left(\langle \tilde{\phi}_n \tilde{\phi}_m \rangle - \langle \tilde{\phi}_n \rangle \langle \tilde{\phi}_m \rangle\right) \\ &\times \left(\delta_{\mu\nu} p(h_\mu) - p(h_\nu) p(h_\mu)\right)\end{aligned}$$

and the information maximization term is

$$\begin{aligned}\partial_n^\mu \partial_m^\nu I(Y,H) = &\left(\left\langle \frac{\langle \tilde{\phi}_n \rangle_{p(\cdot,y)} \langle \tilde{\phi}_m \rangle_{p(\cdot,y)}}{p(y)^2} \right\rangle - \langle \tilde{\phi}_n \rangle \langle \tilde{\phi}_m \rangle\right) \\ &\times \left(\delta_{\mu\nu} p(h_\mu) - p(h_\nu) p(h_\mu)\right)\end{aligned}$$

Angled brackets with no subscript correspond to an average with respect to the steady state distribution, $\langle \cdot \rangle = \sum_{x_i} \cdot\, p(x_i)$. For the second term, we retain the integration variable $y$ for clarity. In sum, the Hessian of the Lagrangian is given by

$$H_{(\mu,n),(\nu,m)} = \left(\langle \tilde{\phi}_n \tilde{\phi}_m \rangle - \beta \left\langle \frac{\langle \tilde{\phi}_n \rangle_{p(\cdot,y)} \langle \tilde{\phi}_m \rangle_{p(\cdot,y)}}{p(y)^2} \right\rangle - (1-\beta)\langle \tilde{\phi}_n \rangle \langle \tilde{\phi}_m \rangle\right)\left(\delta_{\mu\nu} p(h_\mu) - p(h_\nu) p(h_\mu)\right) \tag{23}$$

which lives in $\mathbb{R}^{(N_X \times N_H)\times(N_X \times N_H)}$; we index $H$ with a multi-index $(\mu, n)$. From the form of Eq. 23, we see that $H$ is given by a Kronecker (tensor) product

$$H_{(\mu,n),(\nu,m)} = A^\beta_{nm} \otimes G_{\mu\nu}.$$

Eigenvalues of $H$ are given by products of eigenvalues of $A^\beta$ and $G$, while eigenvectors are given by the tensor product of eigenvectors of $A^\beta$ and $G$. The appearance of unstable directions of the Hessian corresponds to the appearance of negative eigenvalues in its spectrum. As $G$ does not depend on $\beta$, zero-crossings of eigenvalues of $H$ therefore correspond to zero-crossings of eigenvalues in the spectrum of $A^\beta$.

***Equilibrium systems:*** We first study the stability of the matrix $A^\beta_{nm}$ in the case of quasi-equilibrium dynamics. In particular, we consider a Fokker-Planck operator $\mathcal{L}_{\mathrm{FP}}$ of the form

$$\partial_t p(x) = \mathcal{L}_{\mathrm{FP}} p(x) = -\partial_i (f_i(x) p(x)) + D\partial_i^2 p(x). \tag{24}$$

The steady state distribution $\rho_0(x)$ satisfies

$$\mathcal{L}_{\mathrm{FP}} \rho_0(x) = -\partial_i J_i(x) = 0 \tag{25}$$

where $J_i = f_i \rho_0 - D\partial_i \rho_0$ is a flux. Our main assumption in this subsection is that in the steady state, fluxes vanish: $J_i = 0$. In this case, left eigenfunctions $\phi_n$ of $\mathcal{L}_{\mathrm{FP}}$ become *right* eigenfunctions when multiplied by $\rho_0$. To see this, note that

$$\begin{aligned}\mathcal{L}_{\mathrm{FP}}(\rho_0 \phi_n) &= \phi_n \mathcal{L}_{\mathrm{FP}} \rho_0 + \rho_0 \mathcal{L}^\dagger_{\mathrm{FP}} \phi_n - 2\partial_i \phi_n \underbrace{(f_i \rho_0 - D\partial_i \rho_0)}_{J_i} \\ &= \lambda_n \rho_0 \phi_n\end{aligned} \tag{26}$$

where the first term disappears because $\rho_0$ is the steady-state distribution, and the third term disappears because of our above assumption on disappearing fluxes, $J_i = 0$. The following inner product then satisfies

$$\begin{aligned}\langle \phi_n\rho_0, \mathcal{L}^\dagger_{\text{FP}}\phi_m\rangle &= \langle \mathcal{L}_{\text{FP}}\phi_n\rho_0, \phi_m\rangle\\ \lambda_m\langle\phi_n\rho_0,\phi_m\rangle &= \lambda_n\langle\phi_n\rho_0,\phi_m\rangle\\ \rightarrow (\lambda_m-\lambda_n)\langle\phi_n\rho_0,\phi_m\rangle &= 0.\end{aligned}$$

This shows that if $\lambda_n \neq \lambda_m$, the inner product vanishes. Similarly, one can show that the same must be true for $\langle \frac{\rho_n}{\rho_0}, \frac{\rho_m}{\rho_0}\rangle$. These are precisely the types of terms appearing in the Hessian. Consequently, for equilibrium (no flux) systems the Hessian is diagonal. The first term follows directly from the above, while the second is given by

$$\begin{aligned}\sum_y \frac{1}{p(y)}\left(\sum_x p(y,x)\tilde{\phi}_m(x)\right)&\left(\sum_{x'} p(y,x')p_\lambda\tilde{\phi}_n(x')\right)\\ &= \sum_{ij}\sum_y \frac{1}{p(y)}\rho_i(y)\rho_j(y)\mathrm{e}^{(\lambda_i+\lambda_n)\Delta t}\langle\phi_i\rho_0,\phi_n\rangle\\ &\qquad\times \mathrm{e}^{(\lambda_j+\lambda_m)\Delta t}\langle\phi_j\rho_0,\phi_m\rangle\\ &= \sum_{ij}\mathrm{e}^{(\lambda_i+\lambda_n+\lambda_j+\lambda_m)\Delta t}\delta_{ij}\delta_{ni}\delta_{mj}\\ &= \delta_{nm}\mathrm{e}^{(2\lambda_n+2\lambda_m)\Delta t}\end{aligned}$$

The full matrix $A$ appearing in the Hessian then takes the form

$$A_{nm} = \delta_{nm}\mathrm{e}^{2\lambda_n\Delta t}\left(1-\beta\mathrm{e}^{2\lambda_n\Delta t}\right)$$

except for the $n = m = 0$ term, which is $A_{00} = 0$. The eigenvalues are given directly by these diagonal elements. For small $\beta$ these are all positive, and become unstable one after the other at

$$\beta = \mathrm{e}^{-2\lambda_1\Delta t}, \mathrm{e}^{-2\lambda_2\Delta t}, ... \qquad [27]$$

which are increasing in order (remember $\lambda_i \leq 0$). It follows that at the first transition, only $f_1$ becomes non-zero, and hence the encoder takes the form given by Eq. (9). In the equilibrium case, the encoder learns *exclusively* the first eigenfunction, with no correction due to the second eigenfunction.

***General Case:*** We now show that the first component $f_1$ is selected at the first IB transition even when the flux $\vec{J}$ is non-zero. From our assumption on the spectrum of $\mathcal{L}_U$, namely $0 > \mathrm{Re}\lambda_1 > \mathrm{Re}\lambda_2 \gg \mathrm{Re}\lambda_3 ...$ it follows that near the first IB transition the IB loss is given by

$$\mathcal{L}_{\text{IB}} = \begin{pmatrix} f_1\\ f_2\end{pmatrix}^T \begin{pmatrix} a_{11} & a_{12}\\ a_{12} & a_{22}\end{pmatrix}\begin{pmatrix} f_1\\ f_2\end{pmatrix} + \mathcal{O}(f_n^3, \mathrm{e}^{\lambda_3\Delta t})$$

with

$$\begin{aligned}a_{11} &= \mathrm{e}^{2\lambda_1\Delta t}\left(\langle\phi_1^2\rangle - \beta B_{11}\right) = \mathrm{e}^{2\lambda_1\Delta t}\hat{a}_{11}\\ a_{12} &= \mathrm{e}^{(\lambda_1+\lambda_2)\Delta t}\left(\langle\phi_1\phi_2\rangle - \beta B_{12}\right) = \mathrm{e}^{(\lambda_1+\lambda_2)\Delta t}\hat{a}_{12}\\ a_{22} &= \mathrm{e}^{2\lambda_2\Delta t}\left(\langle\phi_2^2\rangle - \beta B_{22}\right) = \mathrm{e}^{2\lambda_2\Delta t}\hat{a}_{22},\end{aligned}$$

where we have introduced the shorthand $B_{ij} = \left\langle \frac{\langle\tilde{\phi}_i\rangle_{p(\cdot,y)}\langle\tilde{\phi}_j\rangle_{p(\cdot,y)}}{p(y)^2}\right\rangle$ and all angled brackets denote averaging with respect to the steady state distribution. The stability of the uniform encoder at $f_n = 0$ is given by the stability of the 2×2 matrix above. The eigenvalues $\omega_i$ and eigenvectors $\vec{v}_i$ of this matrix can be computed explicitly,

$$\begin{aligned}\eta_\pm &= \frac{1}{2}\left(\mathrm{e}^{2\lambda_1\Delta t}\hat{a}_{11} + \mathrm{e}^{2\lambda_2\Delta t}\hat{a}_{22} \pm D\right)\\ \vec{v}_\pm &= \begin{pmatrix}\frac{-1}{2\mathrm{e}^{(\lambda_1+\lambda_2)\Delta t}\hat{a}_{12}}\left(-\mathrm{e}^{2\lambda_1\Delta t}\hat{a}_{11} + \mathrm{e}^{2\lambda_2\Delta t}\hat{a}_{22} \mp D\right)\\ 1\end{pmatrix}\end{aligned}$$

where

$$D = (\mathrm{e}^{4\lambda_1\Delta t}\hat{a}_{11}^2 + 2\mathrm{e}^{2(\lambda_1+\lambda_2)\Delta t}(2\hat{a}_{12}^2 - \hat{a}_{11}\hat{a}_{22}) + \mathrm{e}^{4\lambda_2\Delta t}\hat{a}_{22}^2)^{1/2}.$$

In what follows, we will express quantities in terms of the spectral gap $\Gamma = \lambda_1 - \lambda_2 > 0$. For example, the expression above can be written

$$\begin{aligned}D &= \mathrm{e}^{2\lambda_1\Delta t}\left(\hat{a}_{11}^2 + 2\mathrm{e}^{-2\Gamma\Delta t}(2\hat{a}_{12}^2 - \hat{a}_{11}\hat{a}_{22}) + \mathrm{e}^{-4\Gamma\Delta t}\hat{a}_{22}^2\right)^{1/2}\\ &= \mathrm{e}^{2\lambda_1\Delta t}\left(\hat{a}_{11} + \mathcal{O}(\mathrm{e}^{-2\Gamma\Delta t})\right).\end{aligned}$$

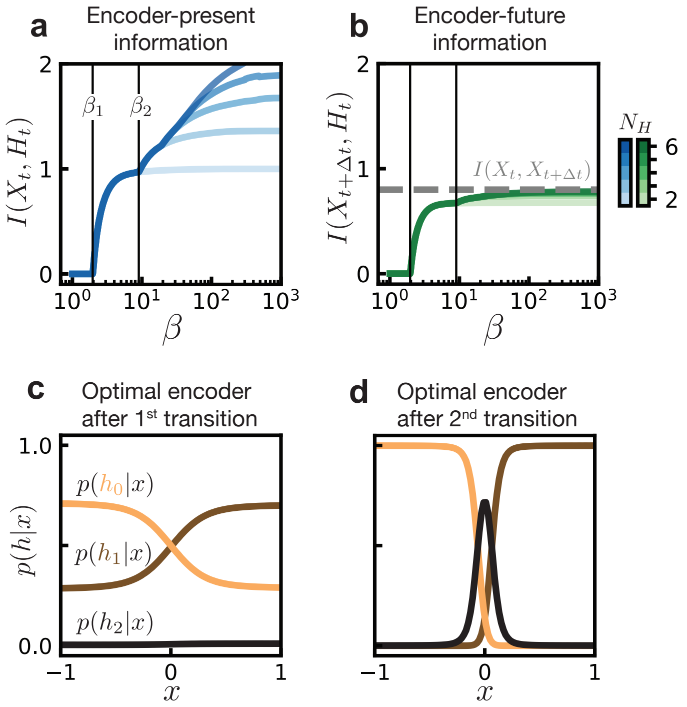


**Fig. S2. Information bottleneck for a Brownian particle in a double well.** (a) Mutual information between the current state $X_t$ and its encoding $H_t$ for various alphabet sizes $N_H$ (color). The first two IB transitions are denoted with black lines and occur at $\beta_1$ and $\beta_2$. (b) Mutual information between the current state's encoding $H_t$ and the future $X_{t+\Delta t}$. The gray dashed line denotes the maximum attainable information, which is the mutual information $I(X_t, X_{t+\Delta t})$. Color denotes encoding alphabet size. Black lines showing IB transitions are shown for reference. (c) Optimal encoder after the first transition ($\beta \gtrsim \beta_1$) with alphabet size $N_H = 3$. (d) Optimal encoder after the second transition ($\beta \gtrsim \beta_2$) with alphabet size $N_H = 3$.

The first eigenvalue to become negative is $\eta_+$. We are interested in the ratio of the corresponding eigenvector components, $f_1/f_2 = \vec{v}_{+,1}/\vec{v}_{+,2}$, which tells us how much the encoder will depend on the first eigenfunction $\phi_1(x)$ compared to the second. This ratio is given by

$$\frac{f_1}{f_2} = \frac{-\mathrm{e}^{(\lambda_1-\lambda_2)\Delta t}}{2\hat{a}_{12}}\left(-\hat{a}_{11} + \mathrm{e}^{2(\lambda_2-\lambda_1)\Delta t}\hat{a}_{22} - D\right) \quad [28]$$

$$= \frac{\mathrm{e}^{\Gamma\Delta t}}{2\hat{a}_{12}}\left(2\hat{a}_{11} + \mathcal{O}(\mathrm{e}^{-2\Gamma\Delta t})\right) \quad [29]$$

$$= \mathrm{e}^{\Gamma\Delta t}\frac{\hat{a}_{11}}{\hat{a}_{12}} + \mathcal{O}(\mathrm{e}^{-2\Gamma\Delta t}). \quad [30]$$

This suggests that we must make one additional assumption, which is that the factor $\hat{a}_{11}/\hat{a}_{12}$ is not small. This is true whenever the flux is small, as

$$\langle\phi_n\phi_m\rangle_{\rho_0} = -\frac{2}{\lambda_n - \lambda_m}\langle(\vec{J}\cdot\nabla\phi_n)\phi_m\rangle_{\rho_0}. \quad [31]$$

Similar to the equilibrium case, we see that at the first transition the encoder depends only on $f_1$, giving Eq. (9). In contrast to the equilibrium case, there may be a small correction due to the second eigenfunction, however this becomes exponentially small for long times $\Delta t$.

Note that in this calculation, $\phi_0(x)$ is constant (following from the assumption of a non-degenerate eigenvalue at 0) so that the $f_0(h)$ factors can be absorbed into $p(h)$. This changes in the case of a degenerate ground state, which corresponds a situation where there are decoupled sectors in which the dynamics evolve independently. Then, each eigenfunction corresponding to the zero eigenvalue is piecewise constant on one of the independent sectors. The optimal encoder Eq. 18 will depend instead on $\phi_0(x)$, which identifies the independent sectors.

### 5. Rate of information decay

We consider the mutual information

$$I(X_t, X_{t+\Delta t}) = \sum_{x,y} p(y|x)p(x)\log\frac{p(y|x)}{p(y)} \quad [32]$$

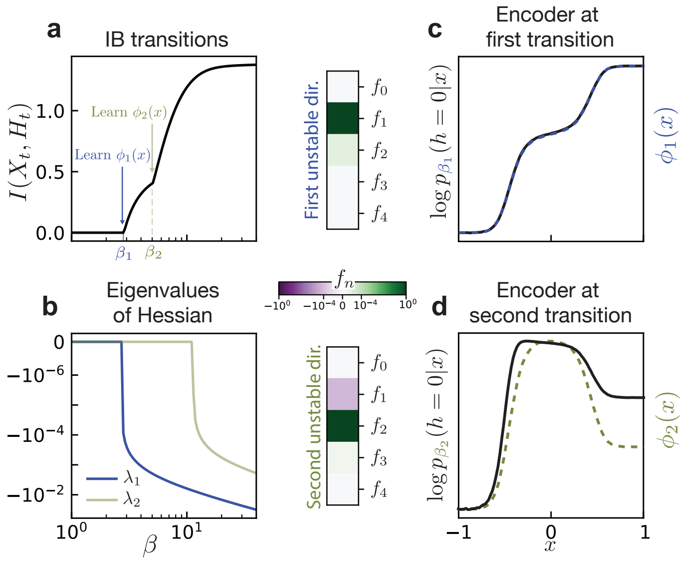


**Fig. S3. IB transitions for a Brownian particle in a triple-well potential.** (a) Information transitions with varying $\beta$ for a particle in a triple-well potential. (b) Eigenvalues of the IB Hessian evaluated at the uniform encoder $p(h|x) = N_H^{-1}$. The appearance of an unstable direction at $\beta_1 \approx 3$ coincides with the first IB transition. The emergence of a second unstable direction doesn't correspond precisely to the IB transition at $\beta_2$ because stability is evaluated at the uniform encoder, but the true optimal encoder at $\beta_2$ is not uniform. (c) The unstable directions are dominated by single components (note the color scale is logarithmic). At the first transition, the logarithm of the encoder is given by the eigenfunction $\phi_1(x)$, up to rescaling ($y$-axis is shown in arbitrary units). (d) Likewise, at the second transition the encoder is given primarily by $\phi_2(x)$.

where $x$ denotes values of the random variable $X_t$ and $y$ the values of $X_{t+\Delta t}$. We further consider dynamics which can be given by a transfer operator $U$ with integral kernel $p(y|x)$ that can be spectrally decomposed as

$$p(y|x) = \sum_n e^{\lambda_n t} \rho_n(y) \phi_n(x), \qquad [33]$$

where $\lambda_0 = 0$, $\phi_0 = \text{const}$ and $\rho_0(y)$ is the steady state distribution. The mutual information can then be written

$$I(X_t, X_{t+\Delta t}) = \sum_{x,y} \sum_n e^{\lambda_n \Delta t} \rho_n(y) \phi_n(x) p(x) \log \left( 1 + \sum_{m>0} e^{\lambda_m \Delta t} \frac{\rho_m(y)}{p(y)} \phi_m(x) \right) \qquad [34]$$

$$= \sum_{n,m>0} e^{(\lambda_n + \lambda_m)\Delta t} \sum_{x,y} \phi_n(x) \phi_m(x) p(x) \frac{\rho_n(y)\rho_m(y)}{p(y)} \qquad [35]$$

$$+ \sum_{n,m,\ell>0} e^{(\lambda_n + \lambda_m + \lambda_\ell)\Delta t} \sum_{x,y} \phi_n(x) \phi_m(x) \phi_\ell(x) p(x) \frac{\rho_n(y)\rho_m(y)\rho_\ell(y)}{p(y)} + \ldots \qquad [36]$$

For long times, the contribution of $\lambda_1$ dominates, with the remaining terms decaying as $\mathrm{e}^{(\lambda_n - \lambda_1)\Delta t}$ for $n \geq 2$. Retaining only the first term, we see

$$I(X_t, X_{t+\Delta t}) = e^{2\lambda_1 \Delta t} \sum_x \phi_1^2(x) p(x) \sum_y \frac{\rho_1(y)^2}{p(y)}. \qquad [37]$$

For short times, other eigenvalues will also contribute to the mutual information (Fig. S4).

### 6. Variational IB

Exactly solving the IB problem in principle requires access to the full distribution $p(y, x)$, where $x$ is the variable to be compressed and $y$ is the relevance variable. One way around this is via so-called Deep Variational IB as introduced in Ref. (13). The key result from (13) is an upper bound on the IB objective

$$\mathcal{L}_{\text{IB}} = I(X; H) - \beta I(Y; H). \qquad [38]$$

To compute the first term, (13) introduces a variational ansatz for the marginal $\hat{p}(h)$. It follows from positivity of the Kullback-Leibler divergence $D_{\text{KL}}(p(h) \| \hat{p}(h))$ that

$$\int dh p(h) \log p(h) \geq \int dh p(h) \log \hat{p}, \qquad [39]$$

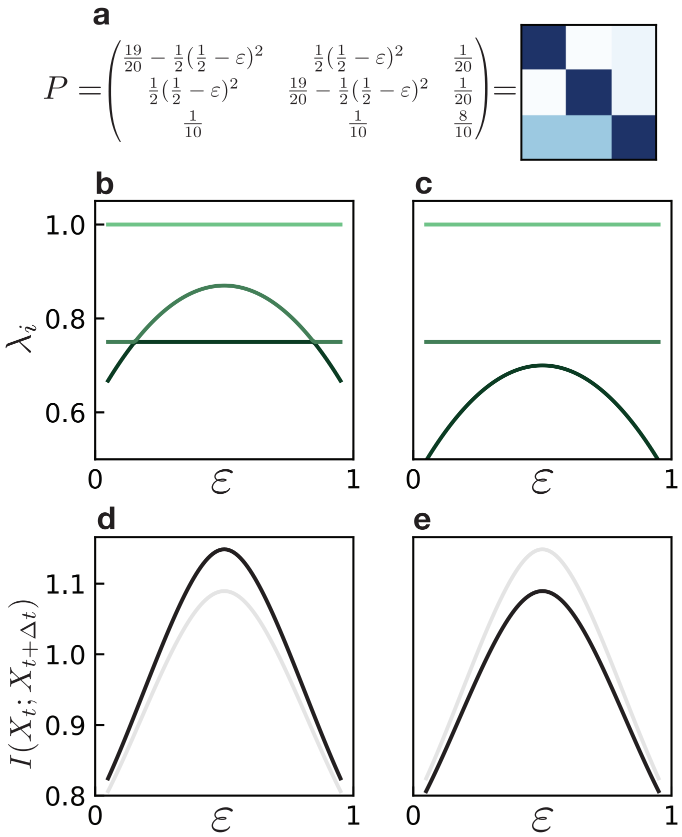


**Fig. S4. Information gain due to an eigenvalue pile-up.** (a) To study the role of a gap closing compared to the pile-up of eigenvalues (beyond the dominant one) in the mutual information $I(X_t; X_{t+\Delta t})$ we build a matrix $P$ with which we can tune the eigenvalues via a parameter $\varepsilon$. (b) Eigenvalues for the matrix shown in (a) with varying parameter values $\varepsilon$; for $\varepsilon \to 0.5$, the spectral gap $\lambda_0 - \lambda_1$ closes. (c) Eigenvalues for the matrix shown in (a), but with all $\frac{1}{2}(\frac{1}{2} - \varepsilon)^2$ terms subtracted with a constant (here $1/8$). Here, the spectral gap $\lambda_0 - \lambda_1$ does not close but there is an accumulation of eigenvalues at $\varepsilon \to 0.5$. (d-e) Mutual information for $\Delta t = 1$ for the matrices in (b-c), respectively, as a function of $\varepsilon$. The curve from the other scenario is shown in gray for comparison. Both exhibit a clear peak in information which differs only slightly in magnitude, showing that the information peak in Fig. 2d-f of the main text is not due to the closing gap alone, but rather due to contributions from subdominant eigenvalues.

and hence

$$I(X,H) \le \int dxdhp(h|x)p(x)\log\frac{p(h|x)}{\hat{p}(h)} \tag{40}$$
$$= \mathbb{E}_x D_{\mathrm{KL}}(p(h|x)\|\hat{p}(h)). \tag{41}$$

If $p(h|x)$ and $\hat{p}(h)$ are chosen properly, the Kullback-Leibler can be expressed analytically which enables gradients to be effectively computed. As in Ref. (13), we take a Gaussian ansatz for $p(h|x)$ and let the marginal $\hat{p}(h)$ be a spherical unit-variance Gaussian. More concretely, encoded variables $H_t$ are sampled from $p(H_t|X_t)$ by computing

$$h_t = f_W(x_t) + \sigma_W(x_t)\eta, \tag{42}$$

where $f_W$ and $\sigma_W$ are deterministic functions modeled by neural networks with parameters (weights) $W$, and $\eta$ is a Gaussian random variable with unit variance.

To bound the entire loss from above, we must bound $I(Y,H)$ from below. We do this using the noise-contrastive estimate of the mutual information introduced in Ref. (14). This recasts the problem as one of distinguishing samples from the distributions $p(y|h)$ and $p(y)$. Given a batch of $B$ pairs $(y,h)$ and one particular value $h_i$, one asks what the probability is that a sample $y_j$ is from $p(y|h_i)$ (is a *positive* sample) and not $p(y)$ (is a *negative* sample). This probability is

$$p(y_j = \mathrm{pos}|h_i) = \frac{\frac{p(y_i|h_i)}{p(y_i)}}{\sum_k^B \frac{p(y_k|h_i)}{p(y_k)}}, \tag{43}$$

where $i$ is the index of the positive sample. The log likelihood of these probabilities,

$$\mathbb{E}[-\log p(y_j = \mathrm{pos}|h_i)], \tag{44}$$

where the expectation is taken over indices $i$ and $j$, is closely related to the mutual information $I(Y,H)$. In the limit of infinite samples $B \to \infty$, this quantity is given by

$$-\int dydh\, p(h)p(y|h)\log p(y_j = \mathrm{pos}|h_i) = \log B - I(Y,H). \tag{45}$$

If one had access to the probabilities $p(y_i = \mathrm{pos}|h)$ appearing in Eq. 44, the mutual information could thus be easily determined. One attempts to estimate these probabilities by introducing a variational ansatz $f(y,h)$ to approximate the density ratio $\frac{p(y|h)}{p(y)}$. Typically this $f$ is represented by a neural network. One can then obtain a bound on the mutual information by minimizing

$$\mathcal{L}_B = \mathbb{E}\left[-\log\frac{f(y_i|h_i)}{\sum_k^B f(y_k|h_i)}\right], \tag{46}$$

from which the InfoNCE estimate of the mutual information can be calculated as

$$I_{\mathrm{NCE}}(Y,H) = \log B - \mathcal{L}_B \le I_{\mathrm{true}}(Y,H). \tag{47}$$

The full objective to be minimized is given by

$$\mathcal{L}_{\mathrm{VIB}} = D_{KL}(p(H_t|X_t)\|\hat{p}(H_t)) - \beta I_{\mathrm{NCE}}(X_{t+\Delta t};H_t), \tag{48}$$

an illustration can be seen in Fig. S5. This loss is evaluated on batches of sample pairs $\{(x_t^{(1)}, x_{t+\Delta t}^{(1)}), (x_t^{(2)}, x_{t+\Delta t}^{(2)}), ...\}$. A minimum is found via stochastic gradient descent. Note that other variational loss functions inspired by the information bottleneck have been derived, for example (15).

The VIB loss Eq. 48 is very similar to the loss function for a time-lagged $\beta$-variational autoencoder ($\beta$-VAE) (16), which have a form $\mathcal{L} = D_{KL} - \mathcal{L}_{\mathrm{rec}}$, where the reconstruction term $\mathcal{L}_{\mathrm{rec}}$ measures the deviation from the true future state $X_{t+\Delta t}$ and the reconstruction from the latent variable $H_t$. In the VIB, this term is replaced by the mutual information $I_{\mathrm{NCE}}$: rather than searching for a latent variable which can reconstruct the While a standard $\beta$-VAE with mean squared error (MSE) loss finds a latent variable that can reconstruct the full state at a time $\Delta t$ in the future, the VIB merely tries to reconstruct the statistics of $X_{t+\Delta t}$ conditioned on $X_t$.

Replacing reconstruction losses with information-theoretical loss functions makes sense in some scenarios, such as chaotic systems. Recent work in this direction has used the Kullback-Leibler divergence as a loss function in place of a $L_2$ loss which generated well-behaved long-term dynamics (17, 18). Other metrics which penalize the number of "false neighbors" in latent space have also shown to improve performance for chaotic systems (19).

The extent to which our findings might apply to time-lagged VAEs with $L_2$ reconstruction losses is an interesting question for future work. In Fig. S13, we observe that time-lagged VAEs learn a similar latent variable as VIB for the simulated fluids dataset. Indeed, several works have noted the apparent similarity between modes learned by variational and regular autoencoders with linear methods such as principal component analysis (PCA) (20–22).

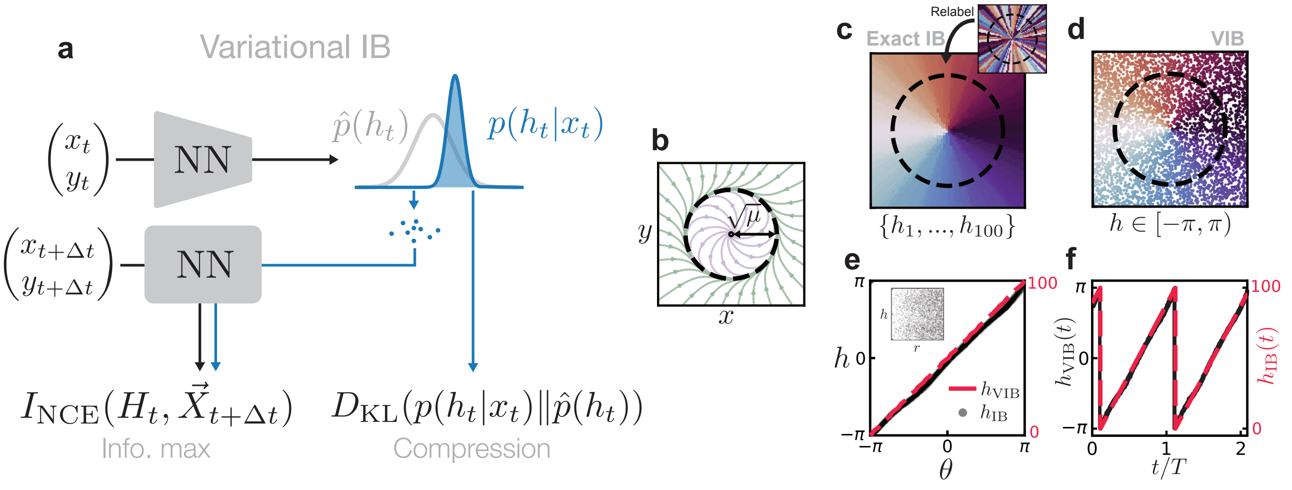


**Fig. S5. Variational IB extends exact IB.** (a) Variational IB optimizes the variational objective directly from samples, in contrast to exact IB which requires an estimation of the full conditional distribution $p(x_{x+\Delta t}|x_t)$. Values of $h_t$ are drawn from a "latent" distribution $p(h|x)$, from which one can estimate the mutual information $I(X_{t+\Delta t}, H_t)$. The compression term is approximated by the Kullback-Leibler divergence between the learned $p(h|x)$ and a variational ansatz for the marginal $\hat{p}(h)$. (b) Phase portrait for a dynamical system above a Hopf bifurcation. (c) Exact IB learns (up to permutations) an encoding $h$ which corresponds to the polar angle coordinate $\theta$. (d) Variational IB similarly learns the correct encoding, directly from samples. (e) Correspondence between IB encodings and the angle coordinate $\theta$. The encoding learned by VIB is independent of $r$ (inset). (f) Time dependence of the encoded variable $h_t$.

**VIB verification for noisy Hopf oscillator.** We now show that VIB learns an encoding consistent with exact IB, i.e. one which depends only on the dominant transfer operator eigenfunctions. We consider the example of a Hopf oscillator given by the dynamical system

$$\dot{r} = r(\mu - r^2), \qquad \dot{\theta} = \omega. \qquad [49]$$

Equation Eq. (49) is the normal form (in polar coordinates) for dynamics near a Hopf bifurcation (23). For $\mu > 0$ this system exhibits a circular limit cycle of radius $\sqrt{\mu}$ (Fig. S5b). In Section 7 we show that this system has infinite purely imaginary eigenvalues $\lambda_n = in\omega$ and that as a result, encoders which exactly encode the angle coordinate, $p(h|r,\theta) \propto \delta(h-\theta)$, are solutions of the IB optimization problem.

The exact IB calculation breaks down for perfectly deterministic dynamics, hence we add a small amount of white noise to the dynamics. While this slightly perturbs the spectral content of the transfer operator, we still find our results to be consistent with the those expected for the deterministic case (see Section 7). As we increase the size of the encoding alphabet $N_H$, the encoder partitions space by finer and finer angular wedges. The learned encoding is invariant under permutations of the encoded symbols. Upon reordering, we find that, for large alphabets $N_H \gg 1$, the encoder indeed approximates $p(h|r,\theta) \propto \delta(h-\theta)$ as expected (Fig. S5c).

Using VIB, we learn a continuous $h_t$ which can be computed directly from samples of the state variable. The encoding $h_t$ learned by VIB closely approximates the angle coordinate (Fig. S5e-f) and is nearly uncorrelated with $r$ (inset). This shows that our mathematical results illustrated for exact IB in Sec. 4 hold also in the approximate framework of VIB.

**VIB for deterministic dynamics.** IB is an inherently probabilistic framework. To handle deterministic dynamics, we make them effectively stochastic by introducing a stochastic sampling scheme. Concretely, rather than taking as our IB relevance variable $Y = X_{t+\Delta t}$, we take $Y = X_{t+\Delta t+\eta}$ where $\eta$ is a random uniformly-distributed time shift. Despite using a different relevance variable, this yields essentially the same optimal encoding as Eq. (9), where the eigenvalue $e^{\lambda_n \Delta t}$ is replaced by $\int d\eta\, p(\eta) \exp(\lambda_n(\Delta t + \eta))$. Crucially, the encoder retains its dependence on the transfer operator eigenfunctions $\phi_n(x)$ as before. In general, this need not be the case: selecting a new relevance variable changes the IB objective and will generically lead to a different encoder. Our choice of stochasticity which we introduce to the dynamics is chosen in such a way to preserve the form of the encoder.

We illustrate this with a prototypical example of deterministic chaotic dynamics, the Lorenz system (24). In the steady state, the state variable of the Lorenz system resides on a chaotic attractor consisting of two "lobes" which encircle unstable fixed points (Fig. S6a). The encoding variable $h$ learned by VIB is the slowest varying function of the state $\vec{x}$, decaying as $\exp(\lambda_1^{\mathrm{Re}} t)$ where $\lambda_1^{\mathrm{Re}}$ is the real part of the first subleading eigenvalue of the transfer operator (here Perron-Frobenius operator) (Fig. S6b). Eigenvalues $\lambda_n$ as well as right eigenvectors $\phi_n(x)$ of the Perron-Frobenius operator were computed numerically using the Ulam method. The correspondence between $\phi_1(x)$ and $h$ disappears as $\beta$ is increased (Fig. S6c-e). The dynamics of $h$ also become notably less "slow", and are instead nearly constant with occasional large jumps (Fig. S6f). This shows that in order for $h$ to be a valid slow variable, the compression term in Eq. Eq. (48) is crucial. These results are consistent with those obtained for exact IB, where we showed that the encoder incorporates the slow modes at low beta (high compression).

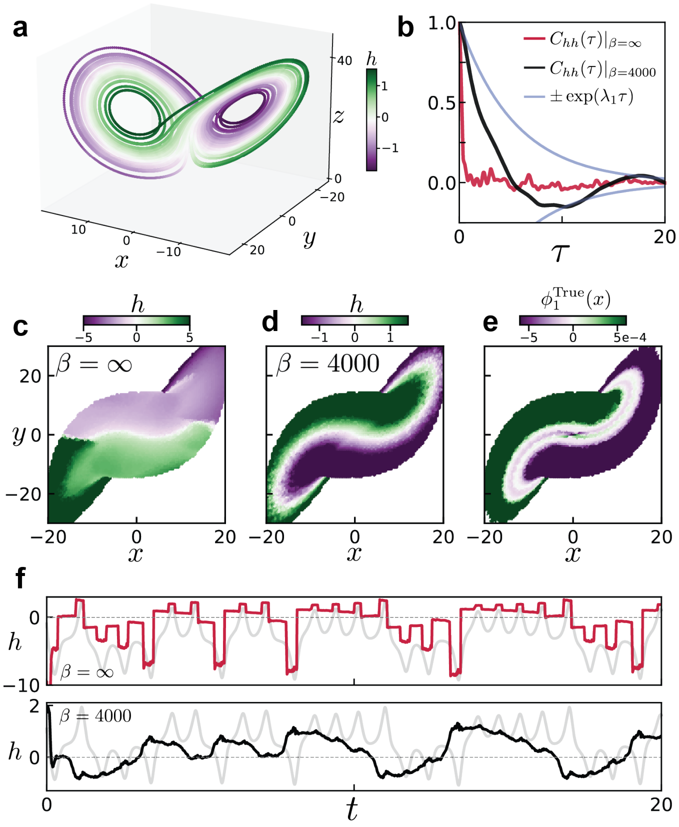


**Fig. S6. Variational IB discovers slow variables.** (a) Variational IB applied to the Lorenz system. Color corresponds the encoding value $h_t$. (b) Time correlation functions of the encoded variable $C_{XY}(\tau) = \langle (X(\tau) - \bar{X})(Y(0) - \bar{Y}) \rangle$ for high compression (black) and no compression (red). Blue shows the decay of the subleading Koopman eigenfunction using numerically-obtained eigenvalues. (c-d) Lorenz attractor projected onto the $x$-$y$ plane, colored by encoding variable. (e) Same projection, colored by the value of the true subleading Koopman eigenfunction. (f) Dynamics of $h_t$ obtained by encoding the state at every time during the true trajectory. Top corresponds to no compression ($\beta = \infty$), bottom corresponds to high compression ($\beta = 4000$).

### 7. Optimal encoding for Hopf normal form dynamics

In Fig. S5 we show that IB learns the angle coordinate when applied to a dynamical system above a Hopf bifurcation. Here, we show that this is expected. We begin by deriving the adjoint transfer operator in polar coordinates, and then continue to solve for its eigenfunctions. In Cartesian coordinates, the equations of motion for the particle are

$$\dot{x} = (\mu - x^2 - y^2)x - \omega y \qquad [50]$$

$$\dot{y} = (\mu - x^2 - y^2)y - \omega x \qquad [51]$$

Expressed in polar coordinates, we have

$$f_x \hat{e}_x = f_x \cos\theta \hat{e}_r - f_x \sin\theta \hat{e}_\theta \qquad [52]$$

$$f_y \hat{e}_y = f_y \sin\theta \hat{e}_r + f_y \cos\theta \hat{e}_\theta \qquad [53]$$

From this we can compute $\mathcal{L}_\mathcal{K}\phi = f_i \partial_i \phi$,

$$f_i \partial_i \phi = (\partial_r \phi)(\mu - r^2)r + \frac{1}{r}(\partial_\theta \phi)\omega r \qquad [54]$$

$$= r(\mu - r^2)(\partial_r \phi) + \omega \partial_\theta \phi \qquad [55]$$

Because this differential equation is separable, we can find eigenfunctions by looking for eigenfunctions that are a function of either $r$ or $\theta$. Recall that for deterministic dynamics, a product of adjoint transfer operator (or *Koopman* operator) eigenfunctions is again an eigenfunction (6). The eigenvalue equation for the radial coordinate is solved in Ref. (7), and in Ref. (25) it is shown that for this system there is no globally valid Koopman decomposition. However we are only interested in a subset of eigenfunctions, in particular those with eigenvalue with real part $\mathrm{Re}\lambda_i \approx 0$, which may not suffice to approximate arbitrary functions of the state variable.

For the angle coordinate alone the situation is much simpler,

$$\omega \partial_\theta \phi(\theta) = \lambda \phi(\theta)$$

which is solved by functions $\phi_n(\theta) = \exp\left(\frac{\lambda_n}{\omega}\theta\right)$. Periodicity requires $\frac{\lambda}{\omega}2\pi = i2\pi n$, which leads to $\lambda = in\omega$. The eigenfunctions and eigenvalues are then

$$\phi_n(\theta) = e^{in\theta}, \qquad \lambda_n = in\omega.$$

The corresponding eigenfunctions of the Perron-Frobenius operator are given by $\rho = \mathrm{e}^{-in\theta}$. We now consider an encoding $p(h|x)$.

$$p(h|r,\theta) \propto \exp\left\{\sum_n^\infty \mathrm{e}^{\lambda_n \Delta t}\phi_n(r,\theta)\int r'dr'd\theta'\rho_n(r',\theta')\log p(r',\theta'|h)\right\} \qquad [56]$$

$$= \exp\left\{\sum_n^\infty \mathrm{e}^{in\omega\Delta t}\int r'dr'd\theta'\mathrm{e}^{in(\theta-\theta')}\log p(r',\theta'|h)\right\} + \mathcal{O}(e^{\mathrm{Re}\lambda_k \Delta t}) \qquad [57]$$

where we retain only the eigenvalues with zero real part; the $\lambda_k$ in the above refer to those eigenvalues with non-zero real part. After neglecting these terms, it can be seen directly that $p(h|r,\theta) = p(h|\theta)$. The coordinates $(r',\theta')$ are used to denote $(r_{t+\Delta t}, \theta_{t+\Delta t})$. The expression can be further simplified by replacing the sum over $n$ with a delta function, which leads to

$$p(h|r,\theta) \propto \exp\left\{\int r'dr'\log p(r',(\theta+\omega\Delta t)|h)\right\}. \qquad [58]$$

We next ask whether an encoder of the form $p(h|\theta) \propto \delta(h-\theta)$ is a solution. Recall that the probability distribution appearing in Eqn. 58 is a distribution over *future* positions, $p(R_{t+\Delta t} = r', \Theta_{t+\Delta t} = \theta + \omega\Delta t|h)$. This distribution can be calculated as

$$p(r_{t+\Delta t}, \theta_{t+\Delta t}|h) = \int r dr d\theta p(r_{t+\Delta t}, \theta_{t+\Delta t}|r_t, \theta_t)p(r, \theta_t|h) \qquad [59]$$

$$\propto \delta((\theta_{t+\Delta t} - \omega\Delta t) - h) \qquad [60]$$

where we used that the two terms in the integrand of the first equation are both delta functions. Plugging this into Eq. (58) shows that $p(h|\theta) \propto \delta(h-\theta)$ is consistent, and hence is an optimal encoding. In the presence of noise, the eigenvectors are perturbed and may gain a dependence on $r$ (see Fig. S7).

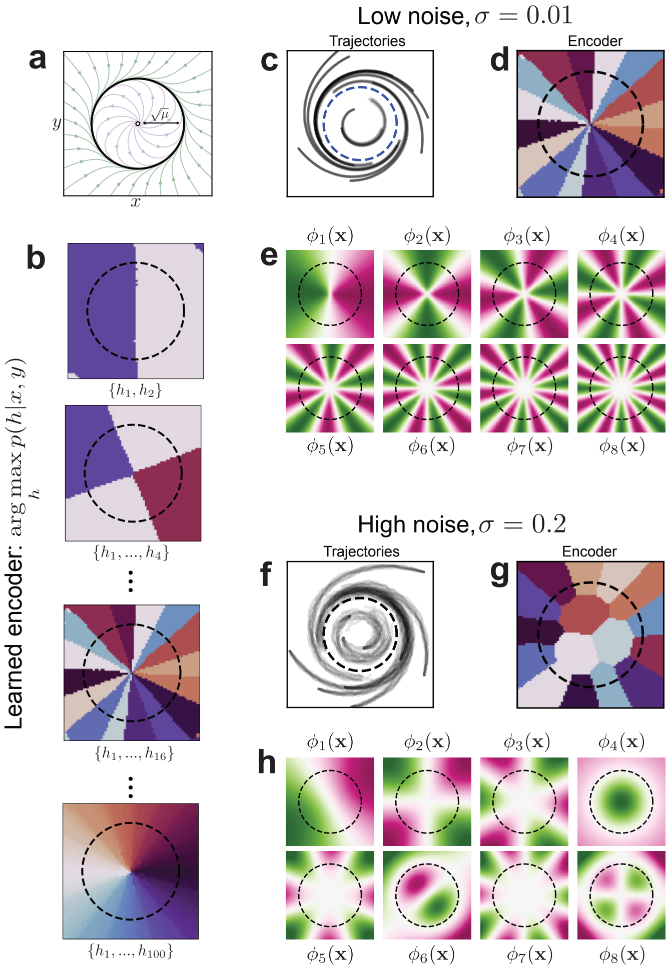


**Fig. S7. Eigenfunctions and IB partitions for the Hopf Oscillator.** (a) Phase portrait for the deterministic Hopf oscillator. (b) Partitions found by IB for high beta but a restricted encoding alphabet $N_H$. (c) Example trajectories of the nearly deterministic Hopf oscillator, with small noise amplitude $\sigma = 0.01$. (d) IB partition of the low noise dynamics for a $N_H = 16$ encoding alphabet (same as appears in (b)). (e) For small noise, the first several eigenfunctions obtained numerically approximate the numerically expected ones of the form $\cos n\theta$. (f) Simulated trajectories of Hopf oscillator with higher noise amplitude $\sigma = 0.2$. (g) IB partition of the low noise dynamics for a $N_H = 16$ encoding alphabet; note the dependence on $r$. (h) The presence of noise changes the eigenfunctions, in particular they depend on $r$.

### 8. VIB for fluid flow around a cylinder

**Eigenfunction computation via dynamic mode decomposition.** Here we provide some additional details on the computation used to generate components of Fig. 5 of the main text. The true Koopman modes were computed using dynamical mode decomposition (DMD) (26–28), also described below in Section 11. DMD attempts to find a finite dimensional approximation of the Koopman operator using $n$ snapshots of the system's state $x \in \mathbb{R}^d$ which are assembled into a data matrix $\mathbf{X} \in \mathbb{R}^{n \times d}$. One then attempts to find a linear evolution operator $K$ which propagates the state forward in time

$$\mathbf{X}_{t+\Delta t} = K\mathbf{X}_t.$$

The approximate Koopman operator is given by the least squares solution $K = \mathbf{X}_{t+\Delta t}\mathbf{X}_t^+$ where $\mathbf{X}_t^+$ denotes the pseudo-inverse of $\mathbf{X}_t$. Approximate Koopman eigenfunctions are given by $\phi_n(x) = x \cdot w_n$ where $w_n$ denotes the $n$th eigenvector of the matrix $K$ and is known as the $n$th DMD mode, denoted by $m^{(n)}$ in the main text.

**Details on gradient analysis.** The fluid flow of a von Karmen vortex street is well approximated by linear dynamics, which can be understood by recognizing that the system is poised just after a Hopf bifurcation so that there is an angle coordinate which rotates with constant angular velocity. This constant rotation can be described by a linear dynamical system. It follows from linearity that eigenfunctions of the adjoint transfer operator are given by linear functions of the state variable,

$$\phi_n[\vec{v}] = \langle \vec{v}(\vec{x}), \vec{m}^{(n)}(\vec{x}) \rangle, \qquad [61]$$

where $\vec{m}^{(n)}$ is the $n$-th "Koopman mode" and angled brackets denote integration over space. To compute these modes and the corresponding transfer operator spectrum, we use dynamic mode decomposition (DMD; see SI Section 8) (26, 27).

We allow VIB to learn a two-dimensional encoding variable $[h_0, h_1]$, so that it can learn the complete first eigenfunction rather than only the real or imaginary part. For the purposes of comparing our learned variable with $\phi_1$, we construct a complex $h = h_0 + ih_1$ out of the two learned components.

To understand which function has been learned by the neural network, we check whether the learned functions $h_i[\vec{v}]$ are of the form Eq. Eq. (61). This can be done by examining gradients of the network with respect to the input field

$$\frac{\partial h}{\partial v_j} = m_j^{(\mathrm{IB})} + g_{\mathrm{res},j}(\vec{v}(\vec{x})) \qquad [62]$$

where we have separated the part of the gradient which is independent of $\vec{v}$ from a residual part which is dependent on $\vec{v}$. Gradients of the true eigenfunctions are given simply by

$$\frac{\partial \phi_n}{\partial v_j} = m_j^{(n)}. \qquad [63]$$

We can then directly compare our latent variables with the true eigenfunctions by comparing their derivatives. If $h$ corresponds to the true eigenfunction, we expect that $\vec{m}^{(\mathrm{IB})}$ is approximately equal to the Koopman mode $\vec{m}^{(1)}$, and that $\vec{g}_{\mathrm{res}}$ is small. While it is unclear how to perform this decomposition in a general setting, we assume that the residual component $g_{\mathrm{res},j}$ averages to zero over an oscillation period of the flow field $\vec{v}$. Then, $m_j^{(\mathrm{IB})} \approx \langle \frac{\partial h}{\partial v_j} \rangle_t$ and $g_{\mathrm{res},j}$ is given by variations about the mean. We see in Fig. S8 that these variations are much smaller than the mean in magnitude, and that they are essentially orthogonal to the mean vector. From this, we conclude that the gradients are given primarily by the constant part $\vec{m}$.

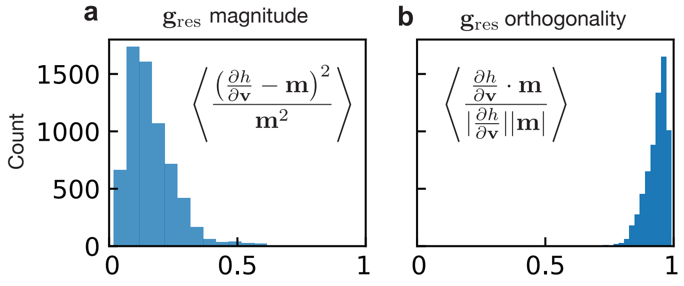


**Fig. S8. VIB gradients for fluid flow are nearly linear.** We compare the linear part of the gradients of the VIB network to the residual which depends on the input field $\vec{v}(\vec{x})$. (a) The residual parts are smaller in magnitude than the linear part $\vec{m}$. Angle brackets denote average over space. (b) The residual parts are nearly orthogonal to the mean $\vec{m}$, as we see that the projection of the full gradient onto the mean is nearly 1.

The learned latent functions vary for different training instances of the neural network. To extract the *average* gradient, we use PCA in an approach similar to that in Ref. (29). The learned functions can have arbitrary sign structure; $h = h_0 + ih_1$ is just as likely to be learned as $h = -h_0 + ih_1$, for example. While in principle the network could learn arbitrary rotations, rather than simply changes in sign, we observe this is not the case. The distribution of gradients forms clusters in the high-dimensional gradient space corresponding to the four possible permutations of sign. PCA picks out the directions separating these clusters, as these are precisely the directions along which the data varies the most. This procedure gives an average gradient, while taking the varying sign structure into account. For reference, the gradient of a single instantiation of the VIB network can be seen in Fig. S13b.

### 9. Cyanobacteria experiments

The experimental parameters used in the cyanobacteria experiments are described in detail in (30). In brief, the authors in (30) control the translation of the KaiA protein with a theophylline riboswitch, allowing them to tune the copy number of KaiA proteins in the bacteria by modulating the concentration of theophylline. The clock state of each individual bacteria is visualized with a fluorescent marker EYFP driven by the *kaiBC* promoter. Colonies are imaged once per hour. The full dataset consists of 5 videos, such as the one shown in Fig. S9, which each contains several colonies. For each video we isolate regions which are filled by bacteria at all times to eliminate the effect of exponential colony growth, which otherwise dominates the VIB results. Each latent trajectory shown in Fig. S9 as well as Fig. 6 in the main text corresponds to the trajectory of one single colony evolving under one of four theophylline conditions. There are 10 trajectories (colonies) in total, coming from 4 experimental conditions (colonies with radius smaller than 64 pixels are not considered). Figure S9 also shows the effect of the choice of time delay, which in the main text we take as $\tau = 3$ hr.

***Synchronization measurements*** We measure the synchronization of the cyanobacterial oscillations using a metric inspired by a locally-coupled Kuramoto model which considers a spatially-varying phase field $\theta(\mathbf{x}, t)$. In terms of this field, an order parameter can be computed as

$$r(t)\mathrm{e}^{i\psi(t)} = \frac{1}{V}\left|\int \mathrm{d}\mathbf{x}\mathrm{e}^{i\theta(\mathbf{x},t)}\right| \qquad [64]$$

where $V$ refers to the volume being integrated over. The value of $r(t)$ is referred to as the synchronization order parameter, while $\psi(t)$ is the average phase.

To calculate the field $\theta(\mathbf{x}, t)$ from an intensity field $I(\mathbf{x}, t)$ we imagine that the intensity field represents one component of the complex phase, for example $I(\mathbf{x}, t) = \sin(\theta(\mathbf{x}, t))$. The other component can be accessed using a time-delay, $\cos(\theta(\mathbf{x}, t)) = \cos(\theta(\mathbf{x}, t + \tau)) = I(\mathbf{x}, t + \tau) =$, where $\tau$ here should be chosen so that the intensity field undergoes one quarter of a full oscillation. As we know the true period of the circadian cycle is 24 hours/frames, we choose a delay of $\tau = 8$ frames. Then, the phase can be computed as

$$\theta(\mathbf{x}, t + \tau) \approx \arctan \frac{I(\mathbf{x}, t)}{I(\mathbf{x}, t + \tau)}.$$

In Fig. S11 we show the results of VIB when applied to a locally-coupled Kuramoto model. Here we learn the same latent features which undergo oscillatory dynamics of varying radius, where the radius corresponds to the synchronization order parameter.

### 10. Simulation Parameters

**Triple well simulations** For the dynamics of the triple well we work directly with the force $F = -\partial_x U$,

$$F(x) = \frac{-1}{200}\left(9375x^5 - 7500x^3 + 1100x - 20\right).$$

The evolution of the Brownian particle's position $x_t$ is given by

$$\mathrm{d}x_t = \frac{F(x_t)}{\gamma}\mathrm{d}t + \sigma\sqrt{\mathrm{d}t}\eta_t \qquad [65]$$

where $\eta_t$ is white noise with unit variance. The noise magnitude $\sigma$ is related to the diffusion constant in Eq. 7 by $D = \frac{\sigma^2}{2}$. We use $\sigma = 1.0$ and $\gamma = 0.2$

Simulations were performed with $N_{\mathrm{init}} = 10^5$ initial conditions, with 300 trajectories generated from each initial condition. The state is evolved for 100 steps at $\mathrm{d}t = 2 \cdot 10^{-3}$. The transfer matrix is approximated by binning the space $x \in [-1, 1]$ with $N_{\mathrm{bins}} = 100$ bins. IB is performed using a time delay $\Delta t = 64$ steps.

**Pitchfork bifurcation simulations** The system was evolved according to the stochastic differential equation Eq. 65 with $F(x) = -\mu x - x^3$, $\gamma = 1$ and $\sigma = 0.1$. For each value of $\mu$, $10^5$ initial conditions were simulated with 2000 trajectories starting at each initial condition. These were evolved for 100 time steps of $\mathrm{d}t = 2 \cdot 10^{-3}$.

**Hopf oscillator simulations** The system was evolved according to Eq. 65 with the force given by Eq.49 in the main text, with $\gamma = 1$ and $\sigma = 10^{-2}$, $\omega = 4.0$ and $\mu = 0.25$. For each value of $\mu$, $10^6$ initial conditions were simulated with 1000 trajectories starting at each initial condition. These were evolved for 50 time steps of $\mathrm{d}t = 2 \cdot 10^{-3}$.

**Lorenz system simulations** The system was evolved according to

$$\dot{x} = \sigma(y - x) \qquad [66]$$
$$\dot{y} = x(\rho - z) - y \qquad [67]$$
$$\dot{z} = xy - \beta z \qquad [68]$$

with $\rho = 28$, $\beta = 8/3$ and $\sigma = 10$. We took $10^3$ initial conditions from which trajectories were simulated for $10^5$ steps with $\mathrm{d}t = 2 \cdot 10^{-3}$. We compute the true eigenfunctions using the GAIO library (31).

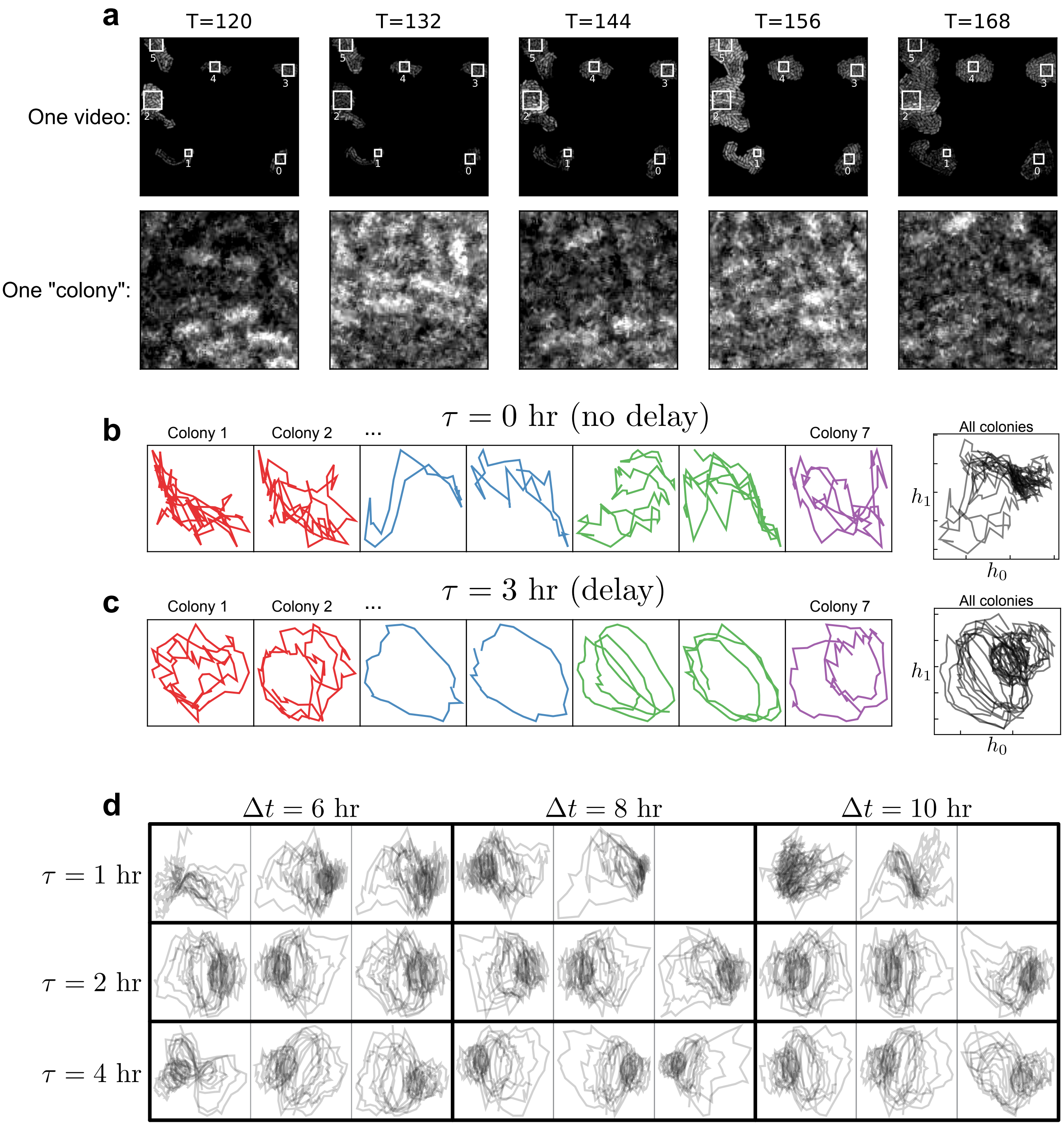


**Fig. S9. Cyanobacteria data and choice of time delay** (a) The cyanobacteria dataset consists of 5 videos, where each has multiple (up to 3) colonies. (Top) time evolution of one video, where the interiors of the colonies are outlined with white boxes. The dataset is composed of all colonies which are greater than 64px in height and width. (Bottom) evolution in time of one selected colony. (b) Learned VIB encodings when no time delay is used, i.e. $\tau = 0$. Left shows the evolution of (a subset of) individual colonies, colored by the original experimental video which they belong to. Note that the axes scale is adjusted for each trajectory so that it fills the plot. Right panel shows all colonies in latent space. (c) Same as above, but with the 3 hr time delay used in the main text. (d) Hyperparameter sweep over different choices of time delay $\tau$ and prediction horizon $\Delta t$. Here each set of three plots was trained with the same parameters but different neural network instantiations to understand how robust these latent spaces are.

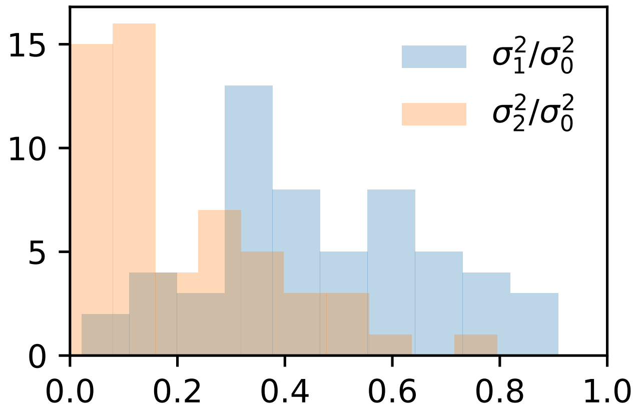


**Fig. S10. Variance along principal component directions** For each DVIB model trained on the cyanobacteria dataset, we compute the principal component decomposition of the resulting point cloud in latent space. Here we show the variance along each principal component for 60 instantiations of the DVIB model. Variances are normalized by the variance in the first principal component direction.

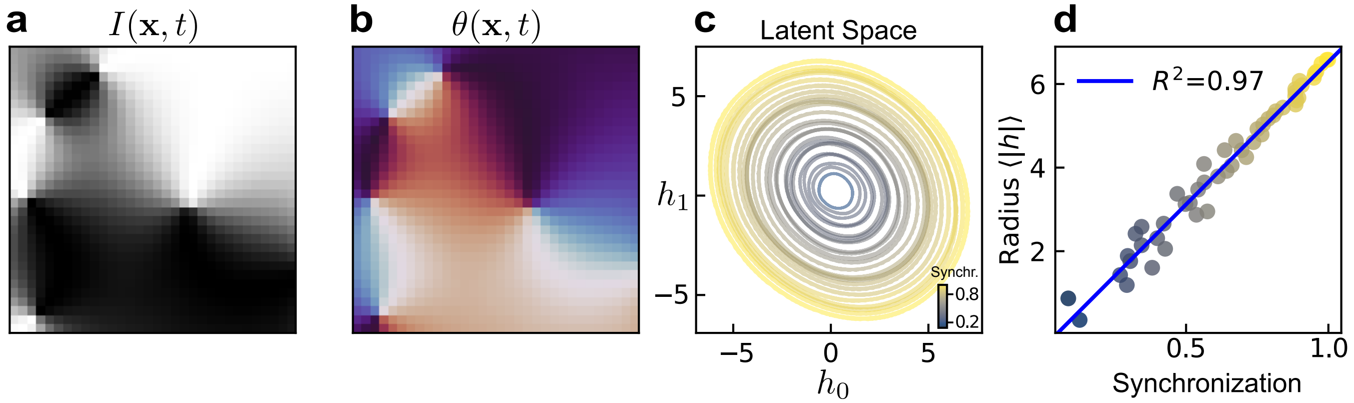


**Fig. S11. VIB applied to simulated Kuramoto model** Inputs are made to mimic the intensity field of the cyanobacteria data. The intensity field (a) is derived from the phase field (b) according to $I(\mathbf{x}, t) = \frac{1}{2}\cos\theta(\mathbf{x}, t) - 1$. (c) DVIB trained with two latent variables learns oscillations. Here each trajectory is colored by its synchronization order parameter. (d) Latent oscillations are very highly correlated with synchronization order parameter.

**Fluid flow simulations** The fluid flow simulations simulations are contained in the "Cylinder in Crossflow" dataset, downloaded from Ref. (32). We take the dataset at Reynolds number 150, and interpolate the velocity field from the unstructured 6569-node mesh to a regular grid of 300 × 150 pixels. The VIB networks are trained with $\beta = 10^7$ and a time buffer $\Delta t$ chosen randomly between $[4, 24]$ (see our comment on time randomization in "VIB for deterministic dynamics" in Section 6).

**Locally-coupled Kuramoto model** In SI Fig. S11 we perform variational IB on a locally-coupled Kuramoto model and find the latent variables learn a synchronization order parameter as in the cyanobacteria data. This model considers a spatially-distributed field of phases which evolves according to

$$\partial_t \theta(\mathbf{x}_i, t) = \omega(\mathbf{x}_i) + J \sum_{j \in \mathcal{N}_i} \sin(\theta(\mathbf{x}_i, t) - \theta(\mathbf{x}_j, t))$$

where $\mathbf{x}_i$ denotes the location of gridpoint $i$ and $\mathcal{N}_i$ denotes the sites neighboring $i$. These dynamics are integrated using the Euler method, where we take $\omega(\mathbf{x}) = \text{const} = 0.05$ for the natural frequencies, and $J = 1.0$.

### 11. Comparison of various dimensionality reduction methods

In this section we compare the performance of variational IB (VIB) to several common data-driven model reduction or inference methods. VIB uses a neural network to identify the relevant variables that are most predictive of the future. This construction learns *dynamically* relevant variables in contrast to methods such as principal component analysis or diffusion maps (in the absence of additional stringent assumptions), and it finds potentially non-linear variables in an agnostic way without requiring a tailored library of non-linearities (such as extended dynamic mode decomposition). In addition to these factors, VIB produces physically well-defined relevant variables – transfer operator eigenfunctions – in contrast to other deep methods such as (variational) autoencoders. In Fig. S12 we compare the low-dimensional latent space trajectories of the reduced models obtained using several of these standard methods to those output by VIB. In the remainder of this section we discuss these methods in detail.

**Principal Component Analysis/Proper Orthogonal Decomposition.** Principal component analysis, or PCA, is a *linear* method that projects the data onto a subspace which accounts for the most variance in the dataset (33). In the dynamical systems literature, PCA is also known as a proper orthogonal decomposition (POD) (34). The starting point for PCA is a collection of *samples* $\mathbf{x}^{(k)}$, where each sample has $N_{\text{feat}}$ different *features*: $\mathbf{x}^{(k)} \in \mathbb{R}^{N_{\text{feat}}}$. Given these samples, one computes a correlation matrix between every pair of features

$$C_{ij} = \frac{1}{N_{\text{samples}}} \sum_{n=1}^{N_{\text{samples}}} (x_i^{(n)} - \bar{x}_i)(x_j^{(n)} - \bar{x}_j).$$

The principal components of the data are eigenvectors of the covariance matrix; the dominant principal component determines the direction of most variation in the dataset. In practice, if many samples are present the correlation matrix is expensive to compute. Writing the dataset as a matrix, $\mathbf{X} \in \mathbb{R}^{N_{\text{samples}} \times N_{\text{feat}}}$, computing the correlation matrix requires evaluating the matrix product $\mathbf{C} = \mathbf{X}^T\mathbf{X}$. Instead, a more computationally efficient way is to compute the singular value decomposition (SVD) of the data matrix,

$$\mathbf{X} = \mathbf{U}\mathbf{\Sigma}\mathbf{V}^T.$$

The principal components are given by columns of $\mathbf{V}$, and the singular values in the matrix $\mathbf{\Sigma}$ are the square root of the variance along each principal component direction. PCA's relation to SVD also means that one can interpret a reconstruction of $\mathbf{X}$ using only $k$ principal components as the optimal rank-$k$ approximation of the dataset, where optimality is here defined with respect to the Frobenius norm.

Compared to VIB, the linearity assumptions underlying PCA can be viewed either as a benefit or a drawback. The benefit is that a linear projection can be intuitive to understand in terms of the original data, and that one can compute this projection quickly. The potential downside is that a linear projection may not be sufficient or appropriate for systems that are highly non-linear. In addition, PCA makes no reference to the dynamics of the system, unlike VIB. An example of where PCA can fail is shown in Fig. 3 of Ref. (35). In words, imagine a particle is hopping in a 2D double-well potential, where the wells are centered at $x \pm 1$. If the vertical extent of the wells is very large ($|y| \gg 0$), then the dominant principal component will be along the $y$-direction, even though the interesting dynamics are the hops between wells, which happens in the $x$-direction.

**Dynamic Mode Decomposition.** Dynamic Mode Decomposition (DMD) starts from the assumption that the system evolves linearly (26, 27). Concretely, if $\mathbf{x}_t \in \mathbb{R}^{n_{\text{feat}}}$ denotes the observed quantity at one point in time, then one assumes dynamics given by

$$\mathbf{x}_{t+1} = \mathbf{A}_{\text{DMD}}\mathbf{x}_t.$$

The matrix $\mathbf{A}$ is a finite-dimensional approximation to the Koopman operator (6).

To find the matrix $\mathbf{A}_{\text{DMD}}$, we start by assembling a collection of samples into a data matrix $\mathbf{X}_t \in \mathbb{R}^{N_{\text{samples}} \times N_{\text{feat}}}$. In addition to this, we assemble a time-shifted matrix $\mathbf{X}_{t+1}$ of the same shape as $\mathbf{X}_t$. In the scenario where we only have one trajectory of duration $T$, each sample may correspond to the system at a measured time point (excluding the final time), so that

$$\mathbf{X}_t = \begin{pmatrix} -\mathbf{x}^{(1)}- \\ -\mathbf{x}^{(2)}- \\ \vdots \\ -\mathbf{x}^{(T-1)}- \end{pmatrix}, \quad \mathbf{X}_{t+1} = \begin{pmatrix} -\mathbf{x}^{(2)}- \\ -\mathbf{x}^{(3)}- \\ \vdots \\ -\mathbf{x}^{(T)}- \end{pmatrix}$$

The matrix $\mathbf{A}_{\text{DMD}}$ is given by the least squares solution

$$\mathbf{A}_{\text{DMD}}^T = \mathbf{X}_t^+\mathbf{X}_{t+1}$$

where $\mathbf{M}^+$ denotes the pseudoinverse of the matrix $\mathbf{M}$. Eigenvectors $\mathbf{w}$ of the matrix $\mathbf{A}_{\text{DMD}}$ are Koopman modes, which correspond to left eigenfunctions of the transfer operator. The evolution of these Koopman modes in time is given by the time-dependent amplitude $a_i(t) = \mathbf{x}_t \cdot \mathbf{w}$, where we take $\mathbf{x}_t$ here to be a single measurement at time $t$.

Similar to PCA, DMD reduces the dimensionality of the dataset by finding a linear projection of the data onto a subspace of the full features space. However, a key difference between the two approaches is that DMD uses the system's dynamics to identify an "optimal" subspace, whereas PCA identifies a subspace based only on the steady-state distribution of the data.

DMD in its original formulation assumes that the Koopman operator linearly evolves the observed state variable. However, the true Koopman operator evolves arbitrary non-linear functions of the state variable forward in time. DMD has therefore been extended to account for non-linear functions through "extended DMD", or eDMD (36). eDMD augments the state vector with non-linear transformations of the state. As an example, a state vector $[-\vec{x}-]$ may be replaced by monomials $[-\vec{x}-, -\vec{x}^2-, -\vec{x}^3-]$ (where here we understand exponentiation as element-wise). Other approaches also exist, such as augmenting the state vector by several time-delayed state vectors (37). Rather than constructing the full $d \times d$ matrix, where $d$ is the dimension of the (possibly augmented) state, one can directly construct a low-rank approximation of $K$ by computing the reduced-rank singular value decomposition (SVD) of the matrix $K$ from the SVDs of $\mathbf{X}_t$ and $\mathbf{X}_{t+\Delta t}$.

Similar to PCA, DMD is primarily limited in its assumption of linear dynamics. In some cases this can be resolved with eDMD, which requires that one identifies a suitable set of non-linear terms to account for potential non-linear eigenfunctions of the Koopman operator. It is unclear how to choose this set of functions in a generic setting, which constitutes the biggest disadvantage to VIB. The features for eDMD must be hand-selected, which in some cases may defeat the purpose of using it as a feature-learning tool. VIB is not subject to this restriction, and can learn relevant features directly from the data.

**Independent Component Analysis.** Independent component analysis (ICA) was originally formulated in the context of blind source separation, where a useful picture is the "cocktail party problem" (38). Imagine you place a set of microphones in a room at a cocktail party; these microphones will record the combination of all the conversations happening at once. The goal of blind source separation is to find a way to, from the recorded signals, isolate the original conversations. Mathematically, ICA finds a solution to the equation

$$\mathbf{x} = \mathbf{A}_{\text{ICA}}\mathbf{s}. \qquad [69]$$

Here, $\mathbf{x}$ denotes the recorded signal, $\mathbf{s}$ denotes the independent sources (conversations), and $\mathbf{A}_{\text{ICA}}$ is the mixing matrix. The only measured quantity is the vector $\mathbf{x}$; both the mixing matrix and the independent sources must be learned.

To find one independent component, we start with an initial guess $y = \mathbf{b}^T\mathbf{x}$ which, for a correct choice of $\mathbf{b}$, should be equal to some component $s_i$. To identify the correct $\mathbf{b}$ we will use the fact that sums of independent variables are more Gaussian than the original variables themselves, which follows from the central limit theorem. By assumption, the observed signals are a linear combination of independent components, so $y$ is also: $y = \mathbf{q}^T\mathbf{s}$. If multiple $q_i$ are non-zero, this will be more non-Gaussian than if only one is non-zero. Thus, $y = s_i$ for the choice of $\mathbf{b}$ for which $y$ is maximally non-Gaussian (38).

Numerically, one optimizes the deviation from Gaussianity using an approximation of the "negentropy" of the distribution of $y$: $\mathcal{J}_{\text{neg}} = S(y_{\text{Gaussian}}) - S(y)$, where $y_{\text{Gaussian}}$ is a normally distribution random variable and $S$ is the Shannon entropy. The intuition for this formulation is that the $\mathcal{J}_{\text{neg}}$ is minimized if $y$ has a unit-Gaussian distribution, so that it serves as a metric for non-Gaussianity. In practice one uses an approximation for $\mathcal{J}_{\text{neg}}$, see Ref. (38) for details. The optimal $\mathbf{b}$ can then be found by doing gradient ascent on this objective function.

On some level, independent component analysis (ICA) is similar to PCA in that it searches for a linear projection of the data onto a subspace of the feature space. In its basic implementation ICA, like PCA, does not incorporate dynamics in contrast to DMD or VIB. The essential difference between PCA and ICA is the assumption of independence of the components. In PCA, unless the data distribution is actually multivariate Gaussian, the components are unlikely to be independent. IB makes no assumptions about the independence of learned encoding variables with each other, only about their level of independence with the original state. We note that ICA can be understood as minimizing the mutual information between encoding variables, $I(H_1; H_2)$, compared to IB's minimization of the mutual information with the original state (38). Which objective is more desirable depends on the system at hand, but in general we do not expect these to lead to the same encodings.

**Time-lagged independent component analysis.** The leveraging of non-Gaussianity in ICA is required due to the fact that, if using whitened data ($\mathbf{z} = \mathbf{V}\mathbf{x}$ so that $\mathbf{z}$ is zero mean ,$\mathbb{E}[\mathbf{z}] = \mathbf{0}$, and unit variance, $\mathbb{E}[\mathbf{z}\mathbf{z}^T] = \mathbf{I}$), then the mixing matrix $\mathbf{A}_{\text{ICA}}$ cannot be estimated if the components $\mathbf{s}$ are normal Gaussian variables: any orthogonal matrix will satisfy Eq. (69). For dynamical systems, however, one can get around this requirement of non-Gaussianity. In particular, one can use a time-lagged variable $\mathbf{z}_{t-\tau} = \mathbf{A}_{\text{ICA}}\mathbf{s}_{t-\tau}$ to compute the mixing matrix from the time-correlation matrix of the signal $\mathbf{z}_t$ (38). In particular, we have

$$\begin{aligned}\mathbb{E}[\mathbf{z}_t\mathbf{z}_{t-\tau}^T] &= \mathbf{A}_{\text{ICA}}\mathbb{E}[\mathbf{s}_t\mathbf{s}_{t-\tau}^T]\mathbf{A}_{\text{ICA}}^T \\ &= \mathbf{A}_{\text{ICA}}\mathbf{D}\mathbf{A}_{\text{ICA}}^T,\end{aligned}$$

where $\mathbf{D}$ is a diagonal matrix. In going to the bottom line, we used the assumption that the components $s_i$ are independent not just instantaneously, but also for a time lag $\tau$. From here we can read off that the matrix $\mathbf{A}_{\text{ICA}}$ is composed of eigenvectors of the correlation matrix of the whitened signal. When written in terms of the original (unwhitened) data $\mathbf{x}_t$ and $\mathbf{x}_{t-\tau}$ it can be shown these eigenvectors are nothing other than the eigenvectors of the transfer matrix $\mathbf{A}_{\text{DMD}}$ (35). Thus, the two methods are equivalent.

**Diffusion maps.** Diffusion maps are a technique which attempts to approximate the Perron-Frobenius operator, or rather the integral kernel $p(x_{t+\Delta t}|x_t)$ (40, 41). Given this approximation, one computes eigenfunctions and uses them as a low-dimensional parameterization of the data ("diffusion coordinates").

This method takes data pairs $\{x_t^{(i)}, x_{t+\Delta t}^{(i)}\}_i$ and approximates the probability of observing these two points via a *kernel* $p(x_t^{(i)}, x_{t+\Delta t}^{(j)}) \approx k(x_t^{(i)}, x_{t+\Delta t}^{(j)})$, where one typically takes a Gaussian Ansatz

$$k_\epsilon(x, y) \propto \exp\left[\frac{-(x-y)^2}{\epsilon}\right].$$

From this, one can assemble the conditional probability distributions into a matrix

$$P_{ij} = p(x_{t+\Delta t}^{(j)}|x_t^{(i)}) = \frac{p(x_t^{(i)}, x_{t+\Delta t}^{(j)})}{p(x_t^{(i)})} = \frac{k(x_t^{(i)}, x_{t+\Delta t}^{(j)})}{\sum_j k(x_t^{(i)}, x_{t+\Delta t}^{(j)})}.$$

This matrix describes the evolution of probability distributions on a *graph* where each node is a data point. In practice, a symmetrized version of $P$ is constructed and the learned eigenvectors are adjusted after the diagonalization (40, 41). To compute the diffusion coordinates of an arbitrary point that wasn't in the original dataset, one inverts the definition of the adjoint transfer operator eigenfunction

$$\phi_i(\vec{x}_{\text{new}}) \approx \frac{1}{\lambda_i} \sum_k P^\dagger_{jk} \phi_i(\vec{x}_k)$$

where $\lambda_i$ is the $i$-th eigenvalue.

Diffusion maps have the advantage, relative to DMD, that they find the full Perron-Frobenius operator and not a linear approximation to it. However, while DMD can isolate the dominant eigenvectors of the operator using reduced-rank SVD, it is less clear how they can be extracted with diffusion maps without first computing the full matrix $P$. We note that VIB, like DMD, also directly learns the dominant modes and does not require estimation of the full transfer operator.

**Deep (Variational) Autoencoders.** Autoencoders (AEs) belong to a class of deep learning methods used for model reduction. Such approaches have successfully been applied in various domains for forecasting the dynamics of complex systems in terms of simpler latent dynamics (42, 43). Autoencoders are composed of an encoder which compresses the observable $\vec{x}$ into a lower-dimensional *latent* variable $\vec{z}$, and a decoder which attempts to reconstruct the original state $\vec{x}$. Variational autoencoders aim to learn a probability distribution over observations $p_\theta(\vec{x}) \approx p(\vec{x})$ from which one can directly sample (44). This is done by assuming that the latent variable is low dimensional, and optimizing the objective

$$\mathcal{L}_{\beta\text{-VAE}} = \mathbb{E}_{q_\phi(z|x)}[-\log p_\theta(x|z)] - \beta D_{\text{KL}}(q(z|x)\|\hat{p}(z))$$

where $q_\phi(z|x)$ is the posterior on the latent variables $z$ and is parameterized by a neural network (encoder) with parameters $\phi$, and $p_\theta(x|z)$ denotes the decoding network with parameters $\theta$. Strictly speaking, we present in the above objective function the $\beta$-VAE loss (16). With $\beta = 1$, which is the case for the original VAE (44), the objective is an upper bound on the log likelihood $-\mathbb{E}[\log p_\theta(x)]$ which is minimized when $p_\theta(x)$ is equal to the true data distribution $p(x)$. The term $\beta$ controls compression as in the IB objective, however it has the opposite effect: small $\beta_{\text{IB}}$ corresponds to high compression, while small $\beta_{\text{VAE}}$ corresponds to low compression.

In cases where one directly computes the probabilities $p_\theta(x|z)$ the first term can be evaluated directly, else it is typically replaced with an $L_2$ loss $\|x - g_\theta(z)\|^2$ (where $g_\theta$ is a deterministic neural network), which is equivalent to assuming a Gaussian Ansatz for $p_\theta$ with a fixed variance.

The VIB loss function is very similar to the $\beta$-VAE loss (45). Rather than attempting to reconstruct the original state $x$, VIB replaces this term with an estimate of the mutual information between the latent variable $z$ and some other relevance variable $y$ (for us, $y = x_{t+\Delta t}$). In contrast to DMD and diffusion maps, neither VIB nor $\beta$-VAEs make any mention of transfer operators and are instead motivated by purely statistical considerations. As we show in the main text, the latent variables learned by VIB correspond to eigenfunctions of the transfer operator. While we cannot claim that that $\beta$-VAEs learn the same thing, some preliminary results in Fig. S13 suggest they may coincide to some degree. In the cyanobacteria dataset, we see that the latent variables learned by a $\beta$-VAE show the same qualitative structure as in VIB, but are less smooth (Fig. S12).

**Other neural networks.** Other deep architectures can also be used for model reduction. For example, Ref. (46) uses recurrent neural networks (RNNs) to learn the evolution of macroscopic variables. To ensure stability and fidelity, the macroscopic variables are periodically "lifted" to the full microscopic state, which is then evolved for several time steps to recalibrate the RNN's hidden state. While interpretability of the latent variables has yet to be explored in such models, we expect the addition of VIB-like objective functions may aid interpretability without harming performance.

As another example, (39) attempts to learn the full Koopman operator using neural networks. This can be thought of as an extension of eDMD, where instead of prescribing the library of nonlinear terms by hand, they can be learned by a neural network. The latent dynamics are then encouraged to be linear by minimizing the deviation between the true future (latent) state $\vec{z}_{t+\Delta t}$ and its linear approximation found by least squares. Similar approaches have also been explored in Refs. (47, 48). As an illustration, we trained one such network on the cyanobacteria which can be seen in Fig. S12. This approach was applied primarily to deterministic systems. We expect that combining their method of encouraging linear latent dynamics together with the VIB objective function may be fruitful and lead to more well-behaved latent variables.

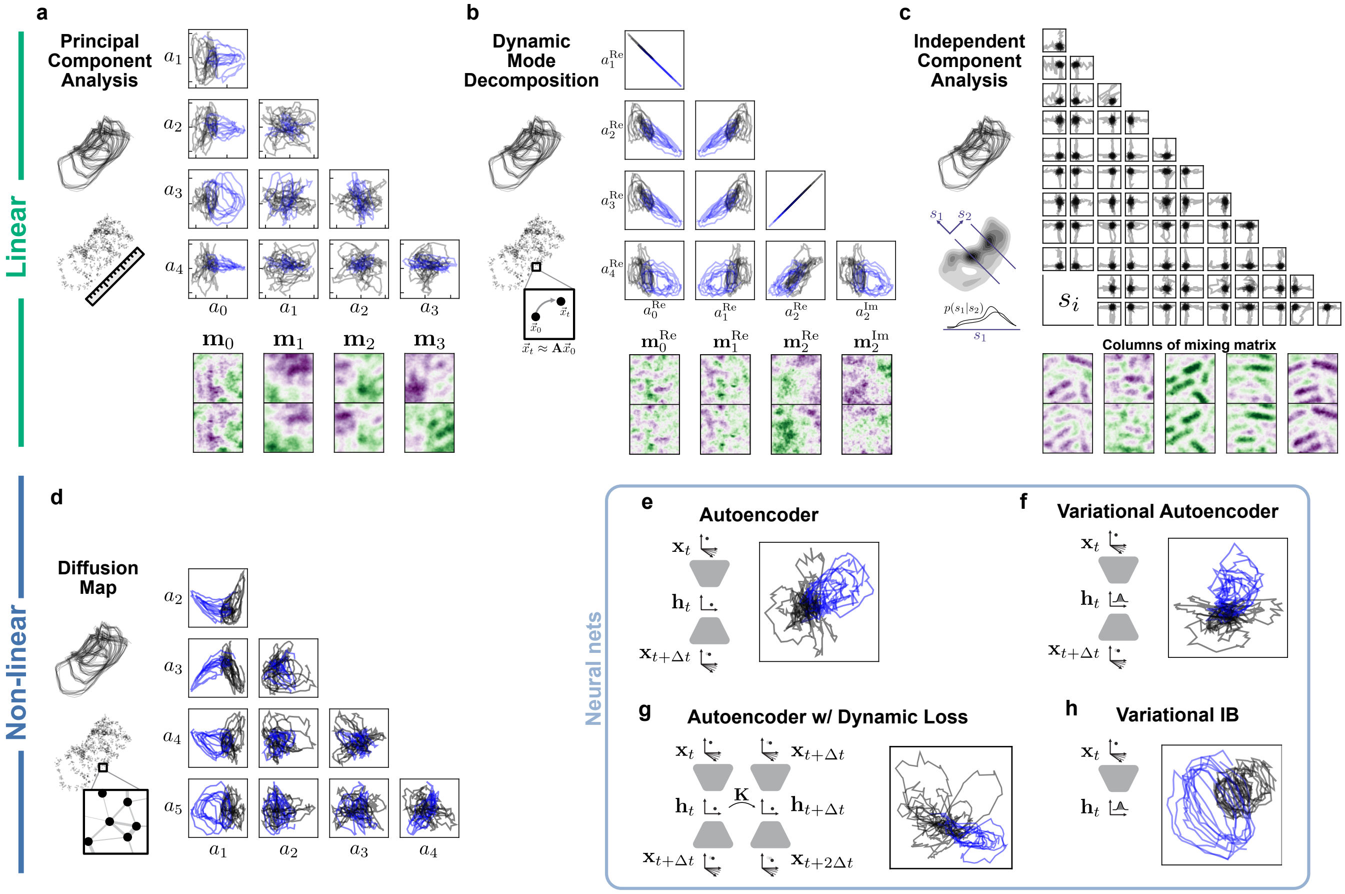


**Fig. S12. Comparison of various dimensionality reduction techniques on cyanobacteria dynamics** Top row illustrates linear methods, bottom row shows non-linear methods. For each method, we plot the trajectories of all cyanobacteria colonies when projected onto the dominant modes (whose definition depends on the particular method). Blue trajectories show the highly synchronized colonies corresponding to high theophylline concentration, black/gray trajectories show the unsynchronized colonies from low theophylline conditions. Unlike the first four methods, deep neural networks (bottom right) directly produce a two-dimensional latent variable and do not provide access to subleading modes. (a) Principal component analysis reduces the dimensionality of data by finding a projection onto the directions of most variance (sketch). Here we show the projection of the cyanobacteria trajectories onto several pairs of principal components, $a_i(t) = \mathbf{x}(t) \cdot \mathbf{m}_i$. Each curve corresponds to the trajectory of one single colony evolving under one of four theophylline conditions; there are 10 trajectories in total, coming from 4 experimental conditions. Below, the first four principal components are shown. Here, the state is composed of two time-lagged images of the cyanobacteria colony; the top half of $\mathbf{m}$ corresponds to the earlier image, the bottom half corresponds to the lagged image. (b) Dynamic mode decomposition is performed on pairs of data points and aims to find a linear evolution operator $\mathbf{A}$ that links them (sketch). (Right) Trajectories of the projection of the state onto several eigenvectors of $\mathbf{A}$. The eigenvectors themselves are shown below (the imaginary parts of the first two modes are uniformly zero). (c) Independent component analysis seeks a linear projection of the data onto statistically independent components $\mathbf{s}_i$. In other words, the distribution $p(s_i|s_j)$ is independent of $s_j$ (sketch). (Right, top) Trajectories of the projection of the state onto several independent components. (Bottom) columns of the mixing matrix; the components $s_i$ describe how the weighting of these columns evolves in time. (d) (Left) Diffusion maps build a graph of the observed state variables with edges weights determined by their distance from one another. (Right) Projection of the trajectories onto the first non-trivial eigenfunctions (because $a_0$ =const) of the inferred transfer operator on the graph. (e) A (time-lagged) autoencoder is a deep neural network architecture which encodes the state $\mathbf{x}_t$ into a low-dimensional latent space. From the encoding one then tries to reconstruct the future state. (Right) Trajectories of the cyanobacterial colonies when encoded into a two-dimensional latent space. (f) A variational autoencoder is similar to a standard autoencoder, where one instead learns a distribution over possible encodings, $p(\mathbf{h}_t|\mathbf{x}_t)$. (Right) Trajectories of the cyanobacterial colonies when encoded into a two-dimensional latent space. (g) Implementation of the network in Ref. (39) which encodes the state using a time-lagged autoencoder with an additional loss term that aims to enforce linearity of the embedded dynamics, $\mathbf{h}_{t+\Delta t} \approx \mathbf{K}\mathbf{h}_t$. (h) Latent trajectories produced by variational IB (VIB).

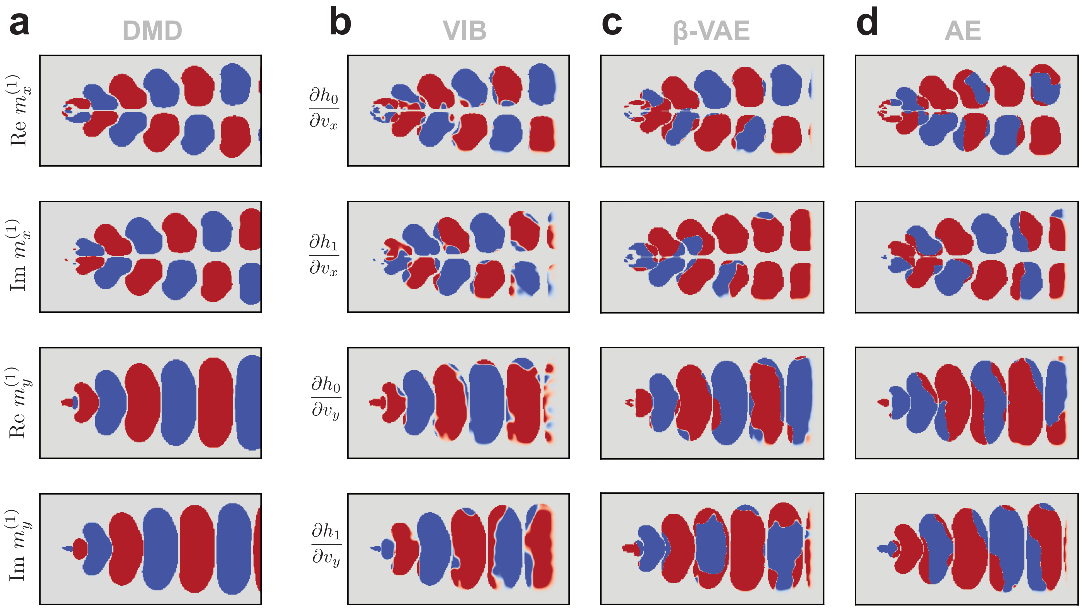


**d** $R^2$

| | $\frac{\partial h_0}{\partial v_x}$ | $\frac{\partial h_1}{\partial v_x}$ | $\frac{\partial h_0}{\partial v_y}$ | $\frac{\partial h_1}{\partial v_y}$ |
|---|---|---|---|---|
| Re $m_x^{(1)}$ | 0.51 | 0.06 | 0.03 | 0.00 |
| Im $m_x^{(1)}$ | 0.01 | 0.43 | 0.00 | 0.00 |
| Re $m_y^{(1)}$ | 0.00 | 0.00 | 0.58 | 0.06 |
| Im $m_y^{(1)}$ | 0.00 | 0.00 | 0.01 | 0.49 |

| | | | |
|---|---|---|---|
| 0.47 | 0.03 | 0.07 | 0.03 |
| 0.00 | 0.25 | 0.05 | 0.00 |
| 0.01 | 0.01 | 0.62 | 0.03 |
| 0.11 | 0.10 | 0.00 | 0.48 |

| | | | |
|---|---|---|---|
| 0.38 | 0.05 | 0.14 | 0.02 |
| 0.00 | 0.37 | 0.00 | 0.11 |
| 0.03 | 0.00 | 0.39 | 0.02 |
| 0.01 | 0.02 | 0.00 | 0.46 |

**Fig. S13. Learned latent variables for fluid flows with various deep neural networks.** To compare latent variables we compare their gradients as in Fig. 5 of the main text, which should correspond to Koopman modes. Here we show the gradients masked by regions in the DMD mode which have large amplitude. (a) True Koopman modes, with real and imaginary parts shown, obtained via DMD. (b) Corresponding modes for a VIB network. Matrix in the bottom row shows the $R^2$ values obtained by regressing the pixel values of the VIB gradients against the different components of the Koopman modes. (c) Corresponding modes for a $\beta$-variational autoencoder (VAE). The VAE has the same encoder structure as the VIB network, while the decoder has an inverted architecture to predict the future state $x_{t+\Delta t}$ from the latent variable $h_t$. Bottom row shows $R^2$ values, analogously to (b). (d) Corresponding modes for an autoencoder.
